# The ENCODE mouse postnatal developmental time course identifies regulatory programs of cell types and cell states

**DOI:** 10.1101/2024.06.12.598567

**Authors:** Elisabeth Rebboah, Narges Rezaie, Brian A. Williams, Annika K. Weimer, Minyi Shi, Xinqiong Yang, Heidi Yahan Liang, Louise A. Dionne, Fairlie Reese, Diane Trout, Jennifer Jou, Ingrid Youngworth, Laura Reinholdt, Samuel Morabito, Michael P. Snyder, Barbara J. Wold, Ali Mortazavi

## Abstract

Postnatal genomic regulation significantly influences tissue and organ maturation but is under-studied relative to existing genomic catalogs of adult tissues or prenatal development in mouse. The ENCODE4 consortium generated the first comprehensive single-nucleus resource of postnatal regulatory events across a diverse set of mouse tissues. The collection spans seven postnatal time points, mirroring human development from childhood to adulthood, and encompasses five core tissues. We identified 30 cell types, further subdivided into 69 subtypes and cell states across adrenal gland, left cerebral cortex, hippocampus, heart, and gastrocnemius muscle. Our annotations cover both known and novel cell differentiation dynamics ranging from early hippocampal neurogenesis to a new sex-specific adrenal gland population during puberty. We used an ensemble Latent Dirichlet Allocation strategy with a curated vocabulary of 2,701 regulatory genes to identify regulatory “topics,” each of which is a gene vector, linked to cell type differentiation, subtype specialization, and transitions between cell states. We find recurrent regulatory topics in tissue-resident macrophages, neural cell types, endothelial cells across multiple tissues, and cycling cells of the adrenal gland and heart. Cell-type-specific topics are enriched in transcription factors and microRNA host genes, while chromatin regulators dominate mitosis topics. Corresponding chromatin accessibility data reveal dynamic and sex-specific regulatory elements, with enriched motifs matching transcription factors in regulatory topics. Together, these analyses identify both tissue-specific and common regulatory programs in postnatal development across multiple tissues through the lens of the factors regulating transcription.

## INTRODUCTION

Mammalian postnatal development is marked by changes in a wide range of biological processes that are coordinated within and between tissues to achieve adult form and function. In both humans and mice, examples include musculoskeletal growth and innervation for locomotion, the neuroendocrine transition at puberty with its sex-specific growth and maturation of both reproductive and non-reproductive tissues, and postnatal brain development necessary for cognition, social behavior and sensory functions. Cell type specializations and cell state transitions underlie these biological processes ^1,2^. Cell types maintain a stable, heritable identity, defined by shared characteristics such as molecular markers, morphology, location, and functional properties. In contrast, cell states represent dynamic variations within a cell type, responding to environmental cues, developmental stages, or physiological changes. These variations involve shifts in gene or protein expression and epigenetic modifications without altering the fundamental cell type ^3,4^. For example, postnatal growth and maturation of skeletal muscle occurs through myofiber growth that includes the addition of new nuclei from differentiating progenitor cells and activity-influenced programming of nuclei within the multinucleate myofibers. These processes lead to distinct type 1 and type 2 fibers with specific contractile properties ^2,5–7^. While myonuclei within muscle cells reflect stable skeletal muscle identity, exercise training can induce cell state transitions between type 1 and type 2 fibers ^2^. To better understand and eventually engineer cell types and transitions between cell states, a first step is the uniform characterization of molecular intermediates such as gene expression and chromatin accessibility at the single-cell level.

Existing single-cell and single-nucleus catalogs primarily capture limited timepoints, focusing on either prenatal development or aging adults. The Tabula Muris Consortium, a widely used resource, recently captured over 350,000 cells in 6 age groups and 23 tissues and organs ^8^, building on their previous *Tabula Muris* catalog of 100,000 cells from 20 organs and tissues using single-cell RNA-seq (scRNA-seq) ^9^. The *Tabula Muris Senis* focused on 1- to 30-month-old mice and identified 155 cell types, averaging around 800 cells per tissue ^8^. Comparative analysis of gene expression across cell types from 3, 18, and 24-month-old mice suggested that certain cell types such as microglia exhibit an intermediate cell state before transitioning to an aged transcriptional profile ^8^. In a focused approach, the systematic dissection of regions in the adult mouse cortex and hippocampus of the Allen Brain Atlas followed by scRNA-seq of 1.3 million cells has produced a comprehensive cell type taxonomy that aligns with the spatial arrangement of the brain ^10^. Although 42 unique subclasses of predominantly GABAergic and glutamatergic neurons were identified, the annotation lacks expected mouse adult stem cells in the brain such as oligodendrocyte precursor cells and neuronal progenitor cells. To provide insights into mouse prenatal development, the ENCODE3 mouse embryo project profiled 12 whole tissues from embryonic day 10.5 to birth using bulk RNA-seq, as well as at the single-nucleus level in forelimb ^11^. This prenatal timecourse of 91,557 total nuclei and 25 cell types revealed dynamic changes in cell type composition and emergence of multiple lineages during skeletal myogenesis in the mouse forelimb. In contrast, our snRNA-seq study spans five core tissues from just after birth to late adulthood at comparable depth to the forelimb time course, pinpointing 99 distinct cell types and states. Our dataset includes an average of around 87,000 nuclei per tissue across 7 timepoints, incorporating 10x Multiome nuclei at two key timepoints.

An ongoing challenge in single-cell resolved transcriptome analysis is to identify and associate groups of genes with meaningful traits. When traits such as sex and age are defined in the metadata, differential expression analysis facilitates the direct comparison of genes enriched in one group compared to another. However, single-cell RNA sequencing notoriously reveals novel cell types and states without clear prior definitions. In such cases, identifying genes associated with these populations presents a significant challenge. While co-expression network analysis has been widely adopted for grouping genes into modules without predefined annotations ^12–14^, it restricts gene membership to a single module. This is problematic because many regulatory genes that define cell type (e.g. transcription factors or cell signaling receptors and transducers) are commonly used recurrently, albeit in differing combinations, across cell types and states. An approach that avoids this limitation starts by identifying ‘cellular programs’, which are distinct sets of genes expressed at specific ratios to one another that can be represented as a vector of weights. A gene can belong to more than one program with different weights or to no program at all. Once the programs are defined, each cell can be scored as expressing a linear combination of the programs. These methods trace their origin to text machine learning used to identify document ‘topics’, so we refer to cellular programs and topics interchangeably. A widely used generative method for topic modeling called Latent Dirichlet Allocation (LDA) can be applied to gene expression data. LDA was originally introduced for population genetics ^15^ and then in natural language processing using machine learning ^16^. More recently, LDA has been repurposed for single-cell RNA-seq to model gene expression by considering genes as words, cells as documents, and latent biological processes as topics ^17,18^. The mixed membership flexibility of LDA aligns with biological reality, where a gene may be repurposed in multiple cellular programs. Analyzing gene weights between topics, which are vectors, facilitates the comparison of attributes and phenotypes associated with a topic, such as dynamic cell types and states, in addition to age and sex.

The core ENCODE4 mouse time course captures postnatal development at key timepoints across cerebral cortex, hippocampus, heart, skeletal muscle, and adrenal glands, encompassing 436,440 total nuclei. We apply LDA using Topyfic with a curated vocabulary of 2,701 regulatory mouse genes ^19^. We recover 82 topics associated with 45 cell types and states including adult stem cells, tissue-resident macrophages, and general proliferation. Using this specific vocabulary allows us to capture cellular programs controlled by transcription factors (TFs) as well as other transcriptional and chromatin regulators such as coactivators, microRNAs, and histone modifiers, and compare them across diverse tissues. Finally, corresponding chromatin accessibility from 10x Multiome at two timepoints ties TFs within our regulatory topics to age-specific and sex-specific cell type- and state-specific regulatory element activity.

## Results

### The ENCODE4 mouse single-nucleus RNA dataset

For the final phase of the ENCODE Consortium, we comprehensively map the mouse polyadenylated RNA transcriptome at the single-nucleus level across 5 coordinated tissues at 7 timepoints in B6/CAST F1 hybrid mice, spanning from postnatal day (PND) 4 to late adulthood (18-20 months) using the Parse Biosciences combinatorial barcoding platform ^20,21^ (Fig. 1a). Complementary genome-wide datasets, including bulk short-read RNA-seq, long-read RNA-seq, microRNA-seq, and chromatin accessibility are also available for matching samples at some or all timepoints (Fig. 1b). Both polyadenylated RNA and chromatin accessibility were measured in the same single nuclei across all five tissues at PND 14 and 2-month timepoints using the 10x Multiome platform ^22^. Notably, this mouse time course mirrors the majority of the human postnatal lifespan, capturing key developmental stages and biological milestones: opening eyes and auditory development, ongoing neurogenesis, synaptogenesis, and myelination especially in the first month of life, adaptation to solid food and social signals after weaning, puberty and sexual maturation by 2 months, and conclusion of the reproductive lifespan by late adulthood.

**Figure 1.**
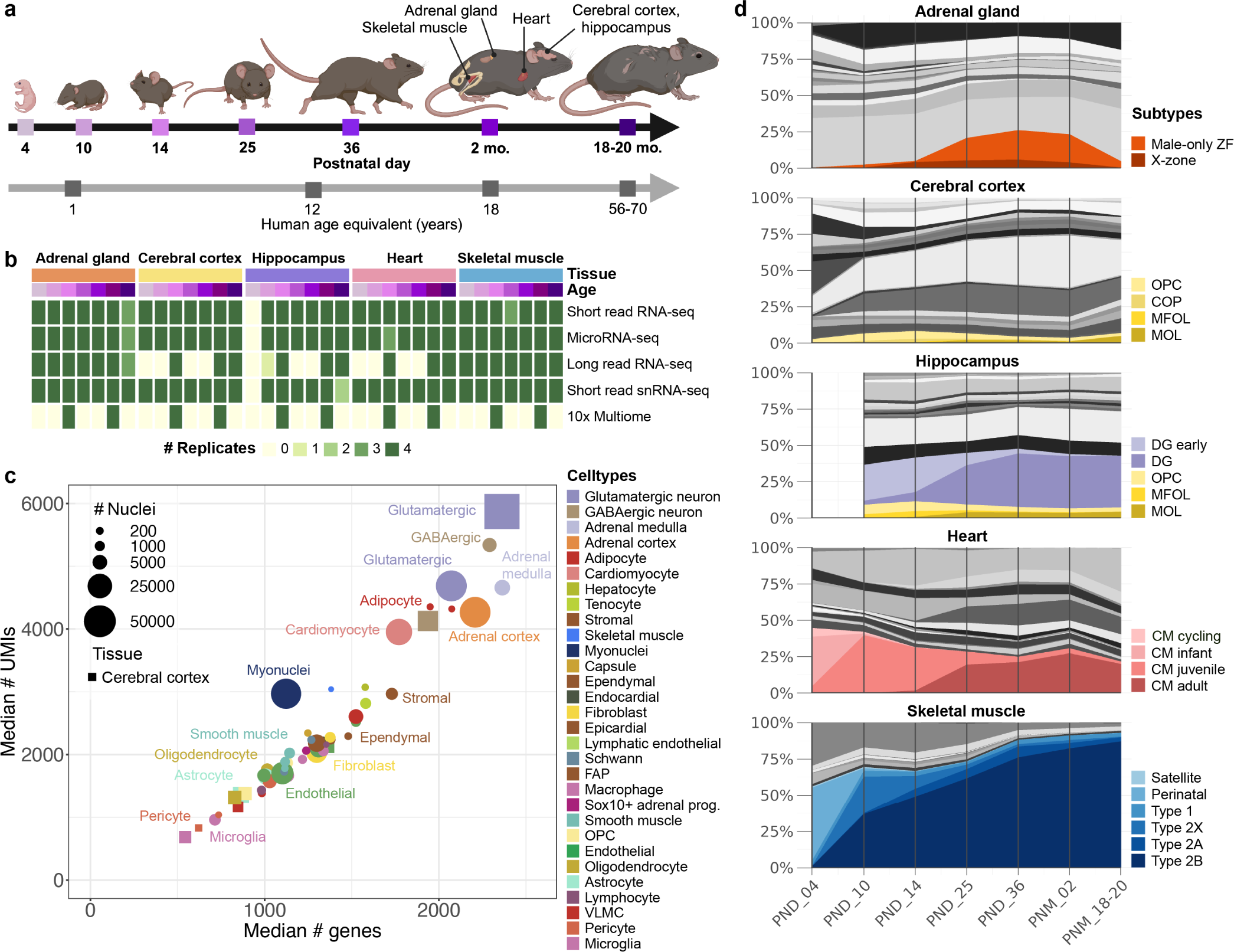
Overview of the ENCODE4 mouse dataset of postnatal development. **a,** Samples from 5 coordinated B6/CAST F1 hybrid mouse tissues were collected at 7 key timepoints from postnatal day 4 to 18-20 months (excluding hippocampus, which was collected from PND 10 onwards). **b,** Overview of the sampled tissues, timepoints, and assays from each tissue in the ENCODE mouse dataset. Most assays have successful experiments in 4 replicates, 2 males and 2 females, per timepoint. 10x Multiome experiments were selectively performed on PND 14 and 2 month timepoints. **c,** Comparison of gene and UMI counts in cell types across all five tissues, with point sizes reflecting the number of nuclei in each cell type within its respective tissue. In common brain cell types, cerebral cortex data points are represented by squares. **d,** Dynamics of subtype composition across postnatal development in all five tissues. Highlighted subtypes are shown in color, while all others are represented in shades of grey (see Fig. S1, S2, S3, S4, S5 full-color versions).

We recovered 83,467 adrenal gland nuclei, 112,118 left cerebral cortex nuclei, 78,168 hippocampus nuclei, 92,808 heart nuclei, and 69,879 skeletal muscle nuclei, collectively expressing 47,707 genes (including protein coding, pseudogene, lncRNA, or microRNA gene biotypes). We annotated each tissue separately for a combined total of 188 clusters, 69 subtypes and states, and 30 major cell types (Fig. S1, S2, S3, S4, S5, Methods). Cells within tissues were clustered, and each cluster was annotated using established marker genes, expert consultations, cluster marker gene identification, literature review, and label transfer from reference datasets where applicable ^10,23–25^(Methods). Our cell type annotations report three hierarchical levels: ‘general cell types’ (e.g. neuron), ‘cell types’ (e.g. ‘GABAergic neurons’), and ‘subtypes’ (e.g. ‘*Pvalb*+’). Every cluster was assigned a single subtype, with larger subtypes spanning multiple Louvain clusters. Cell states were tracked at the subtype level. Evaluation of the number of unique molecular identifier (UMI) counts and genes across cell types reveals reproducible patterns across tissues. Neural cell types such as neurons and adrenal medulla chromaffin cells consistently have more UMIs, and therefore a larger number of detected genes, compared to other cell types such as endothelial and immune cells regardless of the total number of nuclei within each respective cell type (Fig. 1c).

**Figure 2.**
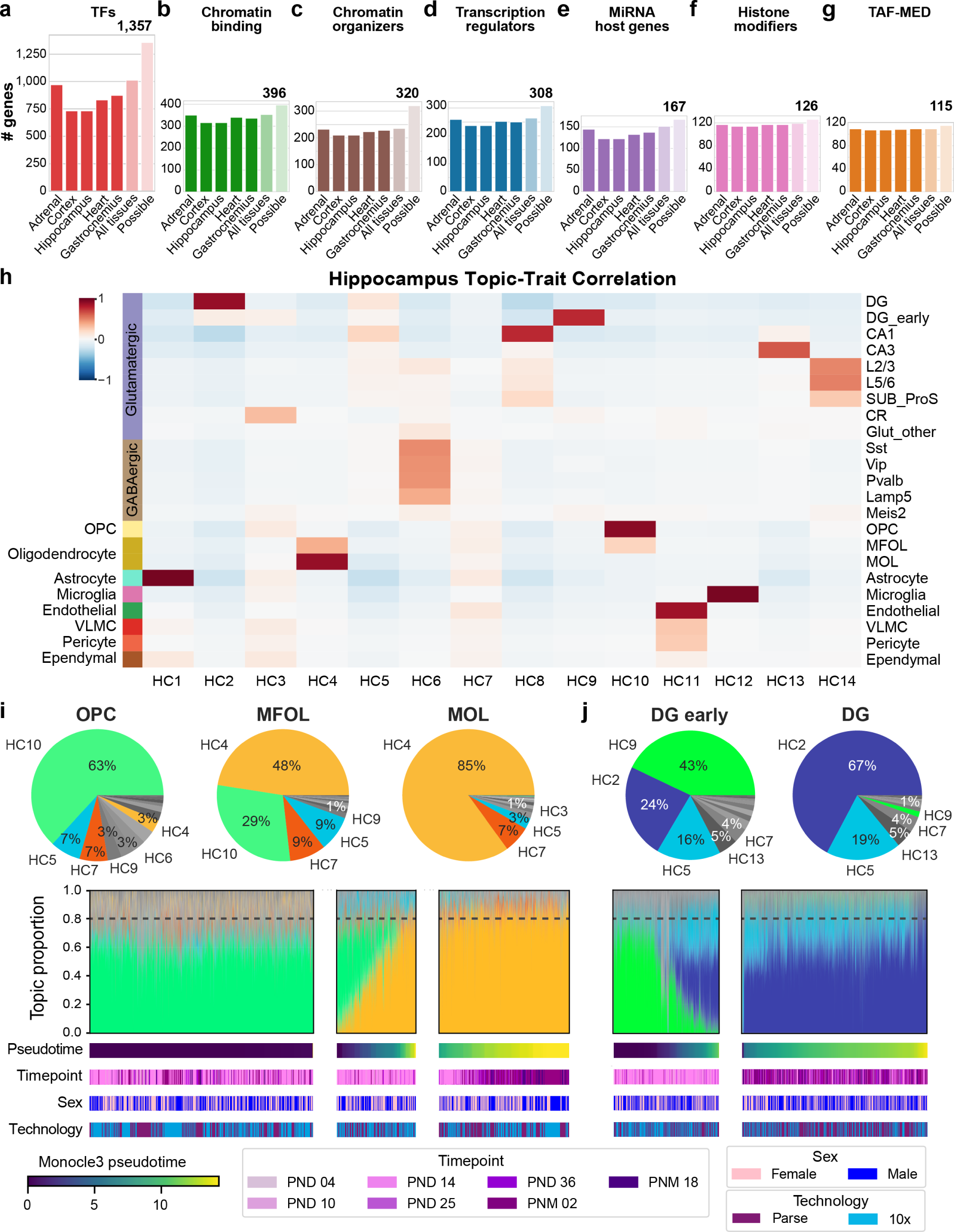
Characterization of hippocampus topics in annotated subtypes. **a,** Number of transcription factors detected at *>* 1 TPM in bulk RNA-seq data in each tissue. Sixth column reports the union of TFs in all tissues, and the last column reports the total number of TFs in our regulatory gene set. **b,** Number of chromatin binding genes, **c,** chromatin organizing genes, **d,** transcription regulators, **e,** host genes representing microRNAs, **f,** histone modifying genes, and **g,** TBP-associated factors and members of the Mediator complex detected in bulk RNA-seq data. **h,** Topic-trait relationship heatmap between 14 hippocampus topics and 10 cell types (23 subtypes). **i,** Proportion of topics in OPC (oligodendrocyte precursor), MFOL (myelin-forming oligodendrocyte), and MOL (mature oligodendrocyte) subtypes summarized in pie charts and displayed as a compressed stacked bar plot (structure plots) for single nuclei ordered by pseudotime. Pseudotime, timepoint, sex, and snRNA-seq barcoding technology are indicated for each nucleus below the structure plots. **j,** Proportion of topics in early DG (dentate gyrus) and DG.

**Figure 3.**
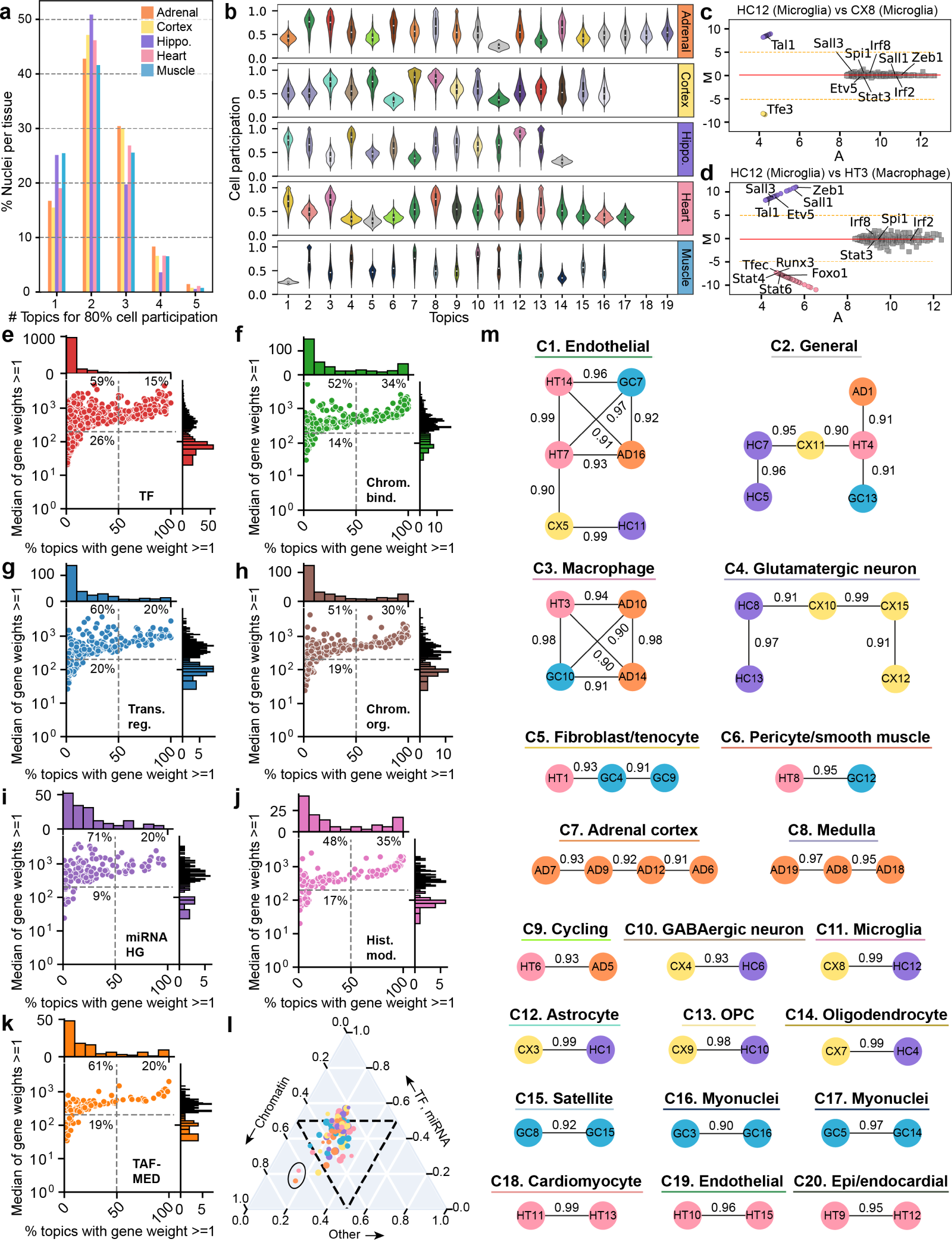
Characterization of topics across diverse tissues. **a,** Comparison of the number of topics required to constitute 80% of cell participation when sorted from the largest to the smallest proportion per nucleus, along with the percentage of nuclei in each category out of the total nuclei per tissue. **b,** Distribution of cell participation in each topic across all five tissues, with violins colored by associated celltype, when possible (see Fig. 1c for color legend). **c,** MA plot comparing HC12 with CX8. X-axis (A) represents average weight of the gene between both topics in the comparison, and y-axis (M) represents log base 2 of the fold change of gene weight between topics. Genes of interest are labeled. **d,** MA plot comparing HC12 with HT3. **e,** Percent of topics containing each gene in the TF biotype vs. median of the gene’s weight across all topics when the gene weight is *>*= 1. Percentages of genes in each quadrant, out of the total number in the biotype, are labeled. Percent of topics containing each gene in each biotype vs. median weight across topics for **f,** chromatin binders, **g,** transcriptional regulators, **h,** chromatin regulators, **i,** microRNA host genes, **j,** histone modifiers, and **k,** TAF-MED complex-associated genes. **l,** Gene biotype simplex with a sector for chromatin (left), encompassing chromatin binders, chromatin regulators, and histone modifiers, a sector for TFs and microRNA host genes (top), and a sector for all other biotypes (right). Topics are color-coded by tissue and scaled by number of genes. **m,** 20 clusters of correlated topics (C1 - C20), filtered to connections *>*= 0.9 cosine similarity. Each node represents a topic, color-coded by tissue, and edges labeled by cosine similarity score calculated on the basis of gene weights between topics.

**Figure 4.**
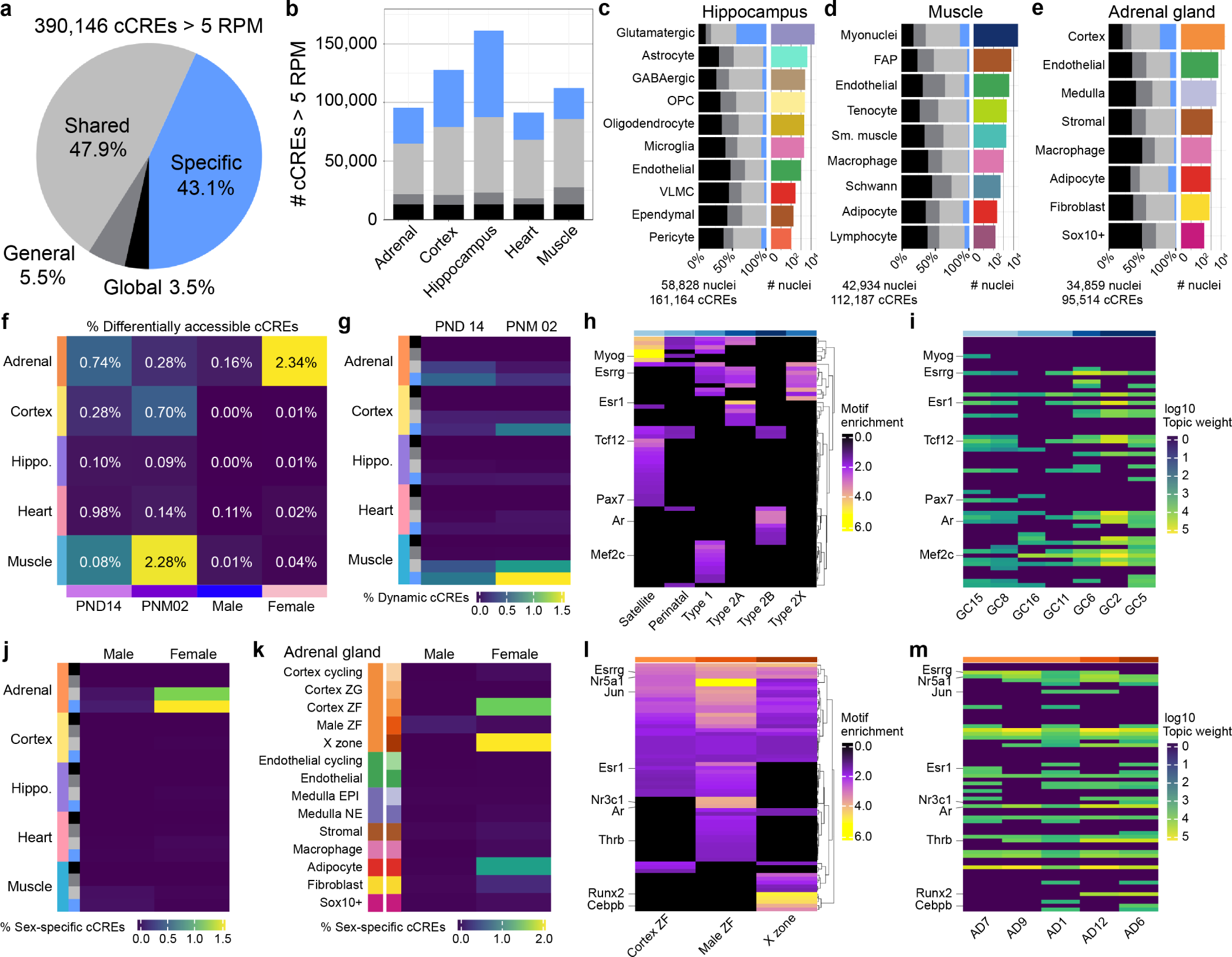
Characterization of celltype-specific candidate cis-regulatory elements and motif enrichment analysis. **a,** 390,146 ENCODE mm10 cCREs filtered by *>* 5 RPM in 10x snATAC-seq data pseudobulked by integrated snRNA-seq clusters. Specific cCREs in blue (168,443) are accessible in only one celltype above 5 RPM across all tissues, shared in grey (186,805) are accessible in more than one celltype within or across tissues, general in dark grey (21,314) are accessible in all major celltypes within a tissue, and global in black (13,584) are accessible in all major celltype across all tissues. **b,** Number of cCRE per specificity category in each tissue. Breakdown of cCRE specificity by percent of cCREs detected in each celltype in **c,** hippocampus, d, gastrocnemius, and **e,** adrenal gland as well as total number of nuclei per celltype. **f,** Percentage of the cCREs detected in each tissue with significant increase in accessibility in each group compared to its counterpart across all tissues. **g,** Overlap of differentially accessible cCREs between timepoints with specificity categories, reported as percent differentially accessible out of total detected in each tissue. **h,** Motif enrichment (adj. p-value *<* 0.05) of expressed TFs (TPM *>* 5 in at least 1 bulk RNA-seq sample) in myonuclear subtype-specific cCREs. **i,** Weight of TFs as ordered in h across topics corresponding to myonuclear subtype-specific subtypes. **j,** Overlap of differentially accessible cCREs between sexes with celltype specificity categories, reported as percent differentially accessible out of total detected in each tissue. **k,** Overlap of sex-specific cCREs with cell-type-specific cCREs, reported as percent differentially accessible out of total detected in each tissue. **l,** Motif enrichment (adj. p-value *<* 0.05) of expressed TFs (TPM *>* 5 in at least 1 bulk RNA-seq sample) in adrenal ZF subtype-specific cCREs. **m,** Weight of TFs as ordered in l across topics corresponding to adrenal ZF subtypes.

### Sex specific layers expand in the adrenal zona fasciculata during puberty before shrinking in late adulthood

Previous studies in B6J mouse adrenal gland characterized the X-zone, a mouse-specific cortical layer situated between the central medulla and the encasing zona fasciculata (ZF) in both male and female mice ^1^. The mouse X-zone and the human fetal zone are both transient cortical layers originating from the fetal stage of development ^1,26^. The human fetal zone disappears rapidly after birth, along with a decrease in steroid secretion, but is functionally similar to the human-specific zona reticularis in adults ^26^. The mouse X-zone becomes detectable by PND 8 and fully emerges as a distinguishable layer by PND 14^1^. In female mice, this layer persists for several weeks during puberty until beginning to regress by PND 32 at the earliest, continuing regression during adulthood. During the first pregnancy, the entire X-zone disappears, while in non-pregnant mice, it undergoes gradual regression before disappearing between 3 and 7 months ^1^. In male mice, the X-zone recedes entirely before PND 40^1^. While the human zona reticularis continues to produce androgens at lower levels after birth, increasing during puberty, mice adrenals lack expression of *Cyp17a1* and thus do not secrete androgens ^27^. Instead, the X-zone is characterized by the expression of 20-alpha-hydroxysteroid dehydrogenase (*Akr1c18*), which has been shown to be induced by estrogen and downregulated by testosterone ^1^. Additionally, *Pik3c2g*, a phosphoinositide 3-kinase involved in cell proliferation, survival, and metabolism is an X-zone marker ^1^. Furthermore, thyroid nuclear hormone receptor beta (*Thrb*) shares X-zone-specific expression with *Akr1c18*. Despite the specificity of these markers, corresponding knockout mouse models lack any X-zone phenotype ^1^. Sex-related factors and other molecules involved in the formation, maintenance, and regression of the X-zone reportedly have no specific expression in the X-zone. Thus, the function of the X-zone remains unclear despite the steroidogenic activity of the fetal adrenal cortex from which it originates.

We identify in males the X-zone counterpart, a large cluster of 8,104 male-specific ZF nuclei that emerges from PND 25 to PND 36 and also regresses in later adulthood (Fig. 1d, S1). Male nuclei make up 95% of the clusters we annotate as male-only ZF, while female nuclei make up 86% of X-zone clusters (4,505 nuclei). We find 303 differentially expressed genes with adjusted p-value *<* 0.01 and log2 fold change (LFC) *>* 1 upregulated in females compared to males in the X-zone and male-specific ZF, including *Xist* and *Tsix* as well as X-zone marker *Pik3c2g* (Methods). *Akr1c18* is not significantly upregulated, but still displays X-zone specific expression (Fig. S1). Ten of the genes upregulated in females are TFs, including *Thrb*, *Runx2*, *Irf8*, and *Nr3c1*. In males compared to females within sex-specific clusters, 666 genes are differentially expressed with adjusted p-value *<* 0.01 and LFC *>* 1, including Y-chromosome linked *Uty* and 35 TFs including *Esrrg* and *Hhex*. Considering these characteristics such as nucleus count, sex specificity, differentially expressed genes, and dynamics mirroring the X-zone in females, we designated the male ZF as a distinct subtype within the broader zona fasciculata in males and females.

### Postnatal neurogenesis and glial maturation in the brain

The hippocampal dentate gyrus (DG) is one of the few brain regions that exhibits postnatal neurogenesis across several mammalian species ^28–30^. In mice and rats, the initial month of postnatal development marks a crucial transitional phase. The most significant maturation shift in the granule cell population occurs between PND 7 and 14^30^. During this period, neuronal progenitor cells (NPCs) expressing doublecortin (Dcx) become localized to the innermost region of the granule cell layer, signifying the establishment of the subgranular zone ^30^. Adult neurogenesis occurs in this specialized niche, from which NPCs eventually migrate to the overlying granule cell layer and become integrated in hippocampal circuitry ^28^. Our data support this narrative, showing that 73% of DG nuclei from PND 10 and PND 14 belong to separate “early DG” clusters whereas 92% of PND 25 and later DG nuclei fall into mature “DG” clusters. Pseudotime ordering from a starting node of cycling nuclei is consistent with real time, distinguishing PND 10 and PND 14 from later timepoints (Methods). Our findings suggest that in later timepoints, the predominant DG cell population is composed of mature *Calb1*+ granule cells; however, approximately a quarter of all our immature *Dcx*+ early DG cells persist into late adulthood (Fig. 1, S3).

Glial maturation is also captured in both the hippocampus and cerebral cortex as a differentiation trajectory from oligodendrocyte precursor cells (OPCs) made up of predominantly early timepoints, though they are present throughout adulthood at lower proportions, to myelin-forming oligodendrocytes (MFOL), to mature oligodendrocytes (MOL) (Fig. 1d, S2, S3). Characterized by the expression of proteoglycan neuron-glial antigen *Cspg4* ^31^, homeodomain transcription factor *Nkx2-2* ^31^, and mitogen *Pdgfra* ^32^, OPCs constitute a highly dynamic and proliferative group of progenitors (Fig. S2, S3). In addition to the primary role of OPCs generating oligodendrocytes in adulthood, OPCs contribute to adaptive myelination and the capacity to regenerate myelin in response to injury or disease ^32^, as well as communicate widely with many of the neural cell types ^33^.

### Cycling and perinatal populations in early postnatal stages of cardiac and skeletal myonuclei

Significant postnatal development occurs in both cardiac and skeletal muscle. In heart, growth is categorized into three phases after birth: hyperplasia until PND 4, rapid hypertrophy between PND 5 and 15, and slow hypertrophy from PND 15 onward ^34^. In our data, proliferating cardiomyocytes marked by expression of *Top2a* and *Mki67* diminish by PND 10, indicating that the first wave of growth is mainly due to cellular division (Fig. 1d). Clustering of ventricular cardiomyocyte nuclei revealed a spectrum of differentiation from infant, juvenile, and adult stages. We find 488 TFs differentially expressed (p. adj *<* 0.01, |LFC*| >* 1) between two or more timepoints in non-cycling ventricular cardiomyocytes, such as genes continually upregulated across postnatal development such as *Foxo3* and retinoid X receptor gamma (*Rxrg*) (Fig. S4, Methods). Several studies have implicated *Foxo3* as a transcriptional regulator of cardiac hypertrophy by inhibiting cardiomyocyte growth and promoting autophagy ^35,36^, potentially responsible in part for the decreased rate of hypertrophy after PND 14. In the mouse embryo, retinoic acid (RA) signaling establishes polarity and promotes the ventricular phenotype in developing cardiomyocytes ^37^, therefore *Rxrg* may also be important in maintaining normal ventricular phenotype in the postnatal state. Cardiomyocyte markers such as *Gata4* and *Mef2* family genes, well-known transcriptional regulators of cardiac genes in infant, juvenile, and adult cardiomyocytes ^38–42^ are expressed throughout development, highlighting the strong regulatory signature of cardiomyocytes at all ages.

As in the brain, skeletal muscle contains adult stem cells, known as satellite cells, that continually replenish myonuclei throughout development and adulthood. As muscles grow, quiescent satellite cells characterized by expression of *Pax7* are activated to become proliferating myoblasts ^43^. Post-mitotic myoblasts align and fuse with each other to form multinucleated myotubes, expressing myogenic regulatory factors (MRFs) including *Myf5*, *Myod1*, and *Myog* ^44,45^. A portion of satellite cells follows an alternative lineage, where they remain unfused and undifferentiated to renew the stem cell pool ^44,45^. Myotubes develop further, undergoing structural organization to become mature myofibers with the ability to perform coordinated contraction and relaxation. Mature skeletal muscle fiber types are identified based on the expression of distinct myosin heavy chain proteins. *Myh7* serves as a marker for slow-twitch type 1 fibers, while *Myh2*, *Myh4*, and *Myh1* are specific to fast-twitch type 2 fibers (2A, 2B, and 2X, respectively) ^6^. Additionally, *Myh3* has classically been linked to embryonic fibers, and *Myh8* to perinatal fibers ^46^. The gastrocnemius, or calf muscle, extends from two heads attached to the femur and in adults is primarily composed of fast-twitch type 2B fibers which run towards the Achilles tendon ^47^. However, fiber type alone provides only a partial understanding of muscle heterogeneity, as the weight of this muscle is sexually dimorphic, with male gastrocnemius weighing 29% more on average than female gastrocnemius at matching timepoints. In our dataset, perinatal myonuclei constitute the majority of myonuclei shortly after birth at PND 4. By PND 10, type 1 myonuclei contribute significantly to the total myonuclei before being surpassed by type 2 fibers, particularly type 2B. However, traces of type 1, as well as type 2A and 2X, persist into adulthood (Fig. 1d, S5). Among 47 single-nucleus clusters, 6 exhibit a notable difference in proportion between males and females, with 5 myonuclei clusters and 1 fibro-adipogenic progenitor cluster showing a difference exceeding 1 standard deviation from the mean (Fig. S5). In addition to tissue-specific cell types, we consistently detect common cell types such as endothelial and immune cells across all our vascularized tissues, maintaining relatively stable proportions. However, their relative proportions in the overall tissue composition varies, with heart tissue having the highest overall counts of endothelial and immune cells (Fig. S1, S2, S3, S4, S5). In summary, our time course effectively captures dynamics of cell types and cell states during postnatal development.

### Topics modeling identifies cellular programs with a core set of regulatory genes

Many genes serve as markers for distinct cell types and states. However, we hypothesize that cellular programs are fundamentally constructed from a core set of genes, including transcription factors (TFs), microRNAs, and chromatin regulators. While a cellular program often controls expression of protein-coding markers that may not be regulators themselves, its core set of regulatory genes governs cell type and state. To study specification of cell types, such as cardiomyocytes, endothelial cells, and microglia, and transitions between cell states, such as transient adrenal cortex zones, granule cell stages, and muscle fiber types, we applied Latent Dirichlet Allocation (LDA) to our annotated snRNA-seq data in each tissue using the Topyfic analysis package ^19^. LDA is a Bayesian model that learns a limited set of hidden topics that can generate the underlying training data ^16^. In the context of single-cell RNA-seq, LDA groups genes into topics and assigns them numerical scores or weights based on their relevance to the topic ^18,19^. By examining the expression patterns of these weighted genes, LDA assigns a participation score to each cell for each topic, ranging from 0 to 1^19^. A participation score of 1 indicates that a cell’s gene expression profile perfectly aligns with the genes associated with that topic ^19^. However, it is rare for a cell to participate in just one topic, as numerous cellular processes are affected by regulatory networks ^48^. Through the analysis of gene weights, LDA enables the comparison of latent traits associated with topics, offering insights into dynamic cell types and states. Topyfic performs LDA 100 times on a normalized ^49^ genes-by-cells matrix and determines consensus topics by clustering all 100 runs ^19,50^. The resulting set of topics represents expression patterns in regulatory genes that define each single cell. These topics can be conceptualized as vectors in gene space, with each weight representing the value in each gene, or dimension. This nuanced approach contrasts with a binary set of marker genes, which merely denotes presence or absence, failing to capture the idea that genes may have multiple roles in different contexts ^51,52^. Overall, the topics approach acknowledges the complexity of cellular programs, recognizing that cells likely participate in multiple programs simultaneously, and underscores the diverse roles that genes may play across various functional contexts.

Our approach to identifying cellular programs involves focusing the LDA vocabulary on genes that we categorize as regulatory. TFs are master regulators of the transcriptome and form the core of cellular programs and gene regulatory networks due to their broad impact on target genes ^53^. Despite their significance, TFs exhibit a wide range of expression patterns across different cell types, often being overshadowed by the expression patterns of their target genes ^54^. In addition to TFs, genes were selected with GO term annotations that impact transcriptional and chromatin regulation such as chromatin binding genes, transcription regulators, chromatin organizing genes, host genes representing microRNAs, histone modifying genes (acetyltransferases, deacetylases, methyltransferases, and demethylases), and TBP-associated factors as well as members of the Mediator complex (TAF-MED) (Methods). Bulk RNA-seq measurements of these genes by regulatory biotype reveals most variation in TF detection at > 1 TPM in at least one bulk sample across tissues (Fig. 2a). Out of 1,357 known TFs in the mouse genome, 1,104 (75%) are detected in one or more tissues, with most in adrenal gland, followed by gastrocnemius and heart, then cortex and hippocampus. Other gene biotypes such as chromatin binding genes, chromatin organizers, and transcription regulators are similarly detected across all tissues (Fig. 2b, c, d). Of the smallest categories (microRNA host genes, TAF-MED, and histone modifiers, Fig. 2e, f, g), the same pattern of adrenal gland, gastrocnemius, heart, and brain regions appears again in the microRNA host gene category, most likely due to the tissue specificity of microRNA expression?. In summary, topics modeling using a curated vocabulary approach aims to extract impactful cellular programs and allows for characterization of regulatory gene biotypes.

### Regulatory gene expression is sufficient to define cell types and cell states

To identify topics specific to each cell type within a tissue, we applied Topyfic on each tissue separately, incorporating batch effect correction between snRNA-seq barcoding platforms ^19,55^. Selecting the appropriate number of topics, denoted as *k*, is a crucial aspect of topic modeling. Topyfic tries different *k* within the range of 5 to 35 for each tissue using 100 random LDA runs per *k* and clusters the resulting topic clusters. If starting with a *k* that is too small, there will be more topic clusters than the starting number *k*, whereas if *k* is too big, it will result in fewer topic clusters than the starting *k*. The optimal *k* is the one that gives as many topic clusters as the starting *k* ^19^. This fine-tuning led to an average of approximately 16 topics per tissue, with the adrenal gland having the highest count at 19, and the hippocampus having the lowest at 14 (Fig. S6, S7, S8, S9, S10, Methods).

Analysis of topic-trait relationships in hippocampal topics indicates that genes crucial for cell type specification are highly weighted in our topics. Topic-trait relationships are analyzed using Spearman correlations to associate specific topics with traits based on cell participation. We observe that hippocampus topic 1 (HC1) corresponds to astrocytes, HC2 to DG granule cells, HC4 to oligodendrocytes, HC6 to inhibitory GABAergic interneurons, HC10 to OPCs, HC11 to endothelial cells, and HC12 to microglia (Fig. 2h). Despite the absence of certain protein-coding genes crucial for cell type-specific functions, such as myelin glycoproteins in oligodendrocytes ^10^, our identified topics exhibit strong correlations with annotated cell types. Developmental progression through the oligodendrocyte lineage is accompanied by topic switching from HC10 in OPCs, to a mix of HC10 and HC4 in intermediate oligodendrocytes (MFOL) to exclusive enrichment of HC4 in mature oligodendrocytes (MOL). Breakdown of cell participation in OPCs and oligodendrocytes shows gradual expansion of HC4 from 3% to 48% to 85%, while HC10 diminishes from 63% in OPCs to 29% in MFOLs during glial differentiation (Fig. 2i). Minor topics HC5 and HC7 remain active throughout differentiation, potentially representing general glial programs that are turned on regardless of subtype. Structure plots are stacked bar plots showing the proportion of topic participation, where each column is a single nucleus grouped by annotated cell type. Ordering of nuclei by pseudotime shows that as cells differentiate, HC10 is gradually replaced by HC4 while minor topics remain constant (Fig. 2i). Notably, topic modeling also captures annotated cell states. HC9 accounts for 43% of the participation of early cells in the DG, while HC2 corresponds to 67% of the participation of mature granule cells (Fig. 2j). Once again, ordering by pseudotime emphasizes topic switching, as HC9 decreases during granule cell maturation. Thus, the expression patterns of regulatory genes alone suffices to define both transcriptional cell types and cell states.

Comparing the number of topics detected per nucleus, we observed that most nuclei in each tissue are effectively characterized by more than one topic, and a median of 2 topics accounts for 80% of cell participation (Fig. 3a). This result supports our hypothesis that cells concurrently run multiple programs, especially during transitional processes of differentiation or maturation ^8^, as evidenced here in hippocampal cell types. Importantly, topics with high cell participation are consistently enriched for specific cell types and states, a trend observed across all tissues (Fig. 3b, S6, S7, S8, S9, S10). Conversely, topics with low participation are typically not associated with any particular cell type (Fig. 3b, shaded gray). At our chosen resolution, all cell types with >1,400 nuclei are captured by at least one topic. In addition to having the highest number of topics compared to other tissues, adrenal gland has the most distinct annotated cell types (10), surpassing other tissues (6, 7, 8, and 8 in cortex, hippocampus, heart, and gastrocnemius, respectively). Interestingly, in both the adrenal gland and heart, a particular topic consistently showed enrichment in cycling cells, irrespective of their cell type of origin (Fig. S6, S9).

### Tissue-specific signals in microglia and macrophage topics

Immune cells are represented by topics with high cell participation across all five tissues. In cortex and hippocampus, topics CX8 and HC12 are associated with microglia, while AD14, HT3, and GC10 correspond to resident macrophages in the adrenal gland, heart, and gastrocnemius, respectively (Fig. 3a). Microglia, the brain’s resident immune cells, originate from progenitors formed during the first wave of primitive hematopoiesis around embryonic day (E) 7.5^56,57^. They migrate to the developing central nervous system (CNS) through the bloodstream, typically around E9.5 in mice ^58^. After prenatal establishment in the CNS, microglia undergo proliferation and expansion, reaching their peak two weeks after birth and sustained through low proliferation levels into adulthood ^58^. The second wave of hematopoiesis gives rise to yolk sac macrophages, a portion of which expand and differentiate into tissue-resident macrophages by E9.5^56^. While previous studies have compared the gene expression profiles of macrophages and microglia derived from adult human brain and blood in culture ^59,60^, as well as infiltrating macrophages and microglia in adult rat brain ^61^, our data and analysis leverage multiple coordinated tissues from the same individual mice.

An MA plot of gene weights for the microglial topics in hippocampus (HC12) vs cortex (CX8) reveals very similar topic compositions, aligning with our expectations (Fig. 3c). Very few genes have an absolute log ratio (M) value *>* 5 (47 in hippocampus, 8 in cortex) (Fig. 3c), none of which have been implicated in regional microglial signatures. Genes involved in microglia polarization (e.g., *Irf8* ^62^ and *Stat3* ^63^), activation and inflammatory response (e.g., *Spi1* ^64^ and *Irf2* ^65^), and establishment of microglia identity and immune response (e.g. *Sall1* ^66^, *Sall3* ^67^, *Etv5* ^68^, and *Zeb1* ^69^ all have high mean average (A) values in both cortex and hippocampus microglia topics. Thus, regulatory topics assign similar weights for genes from identical cell types in different tissues when trained independently.

By contrast, comparison of hippocampus microglia topic HC12 and heart macrophage topic HT3 reveals 165 genes with |M*| >* 5 (67 in hippocampus, 98 in heart). Microglia-specific genes such as *Sall1*, *Sall3*, *Etv5*, and *Zeb1* are more highly weighted in hippocampus, whereas genes involved in macrophage differentiation, polarization, and inflammatory pathway signaling such as *Runx3* ^70^, *Foxo1* ^71,72^, and *Tfec* ^73,74^ exhibit higher weights in heart (Fig. 3d). Interestingly, *Tfec* expression has been shown to be activated by Stat6, another heart-specific macrophage TF in our comparison, which transduces IL-4 signals and binds to the promoter of *Tfec* ^74^ (Fig. 3d). Additionally, *Foxo1* expression has been linked to cardiac fibrosis following macrophage activation ^75^. Due to their similar weights across topics in both tissues, *Spi1*, *Irf2*, *Irf8*, and *Stat3* may belong to a common transcriptional signature of shared immune functions between postnatal microglia and macrophages.

### Mitosis topics are driven by chromatin regulators

We then asked whether particular classes of regulatory genes were found in most topics or were more specific to a subset of topics. We calculated the percentage of topics where a gene surpasses a minimal weight threshold of 1 compared to the median of its weight across all topics (Fig. 3e-k). Notably, 30% or more of genes classified as chromatin regulators (Fig. 3f, h, j) occupy the upper right quadrant, indicating they are highly weighted in most topics. In contrast, transcription factors, transcription regulators, microRNA host genes, and the TAF and Mediator complex family of genes exhibit a different pattern, with 20% or less highly weighted in most topics (Fig. 3e, g, i, k). TFs are mostly either highly weighted and topic-specific (59%, upper left quadrant) or specific with lower weights (26%, lower left quadrant). A simplified analysis of gene biotype enrichment within topics revealed two topics (HT6 and AD5) highly enriched for chromatin regulators compared to TFs and microRNA host genes (Fig. 3j, Methods). Interestingly, these topics correspond to our cycling topics, primarily influenced by a proliferative state rather than their cell type of origin (Fig. S6, S9). Our results suggest that cellular programs essential for mitosis, particularly those governing chromatin condensation and structure, are primarily orchestrated by chromatin regulators. In contrast, programs driven by transcription factors play a lesser role in directing a proliferative cell state.

### Topics in shared cell types from diverse tissues cluster together

We can use cosine similarity, which measures the angle between two topics in gene space, to evaluate differences in relative gene weights. It is similar to other correlation methods, where 0 indicates low concordance between topics and 1 represents high concordance for positive gene weights. By computing the cosine similarity for each pair of topics among the 82 total topics, and subsequently filtering clusters for those with a cosine similarity above 0.9, we identified 20 distinct clusters of topics (Fig. 3m, Methods). As expected, cycling topics HT6 and AD5 are highly correlated with a cosine similarity of 0.93, along with a large cluster of endothelial topics across all five tissues (C9 and C1, respectively, Fig. 3m). Topics representing common cell types across brain regions cluster in C4 (glutamatergic neurons), C10 (GABAergic interneurons), C11 (microglia), C12 (astrocytes), C13 (OPC), and C14 (oligodendrocytes). Interestingly, the macrophage cluster C3 is distinct from the microglia cluster C11. As observed in comparing HC12 and HT3 (cosine similarity 0.83, Fig. 3d), tissue-specific signatures in macrophages and microglia likely drive the differences in gene weights between microglia and macrophage topics. C1 includes two cardiac heart topics, while C19 and C20 represent additional signatures in cardiac endothelial and endocardial cells, distinct from the general endothelial signature shared across all five tissues. In summary, the regulatory topics capture core cellular programs that can be compared across tissues with related cell types.

### Characterizing cell type specificity in candidate cis-regulatory elements

TFs regulate expression of target genes by binding to cis-regulatory elements (CREs) in open chromatin ^54^. The landscape of open chromatin, measured using single nucleus ATAC-seq, provides insight into accessible regulatory elements at the single-cell level. We leveraged the ENCODE registry of candidate cis-regulatory elements (cCREs) in mouse derived from chromatin accessibility, histone modifications, and DNA affinity purification sequencing ^76^ to score our snATAC-seq data across a consistently-defined set of chromatin regions. These elements play crucial roles in gene regulation by providing binding sites for transcription factors and influencing chromatin accessibility ^76^. Around 43% of these regions are classified as candidate distal enhancers by H3K27ac and DNase I hypersensitivity, 12% as proximal enhancers, and 31% were determined by chromatin accessibility data alone (Fig. S11). Accessibility across the full set of 926,843 cCREs was scored in pseudobulk snATAC nuclei using the integrated clusters from snRNA-seq analysis. The cCREs *>*5 RPM in at least one pseudobulk cluster per annotated cell type (390,146 total across our tissues) were classified as specific, shared, general, or global by mapping each cluster to its annotated cell type. We categorized cCREs accessible in only one cell type as ‘specific’, those accessible in more than one cell type within or across tissues as ‘shared’, those accessible in all major cell types within a tissue as ‘general’, and cCREs accessible in all major cell types across all tissues as ‘global’. Most cCREs are either specific to one cell type (43.1%) or shared (47.9%), with only 9% classified as general or global (Fig. 4a). The cell-type-specific landscape of accessible regulatory elements, particularly enhancers, sets the stage for transcription factors to bind and dynamically control gene expression during postnatal development.

Tissue-specific analysis reveals the most cell type-specific elements in cerebral cortex and hippocampus, driven by robust neuronal signatures, with heart displaying the least cell type specificity (Fig. 4b). Indeed, breakdown by cell type in the hippocampus emphasizes glutamatergic neurons as the most specific, and to a lesser extent microglia and pericytes (Fig. 4c). In other tissues, the major cell type also exhibits a robust chromatin signature, such as myonuclei in the gastrocnemius and cortical cells in the adrenal gland (Fig. 4d, e). To further explore the dynamics and sex specificity of the chromatin landscape, which likely contribute to variations between certain cell types, differential accessibility analyses were conducted between timepoints and sexes in accessible cCREs. The largest proportion of differentially accessible cCREs between PND 14 and 2 months are detected in gastrocnemius tissue, while most sex-differential cCREs are detected in adrenal gland (Fig. 4f). This is consistent with biological processes in the major cell types of these tissues; myonuclei in the gastrocnemius are transitioning to their mature fiber type, and the X-zone is emerging in the adrenal zona fasciculata during puberty, emphasizing the dynamic nature of chromatin accessibility during crucial postnatal stages.

### Regulatory motifs are enriched in cell-type-specific cCREs

Although most perinatal myonuclei disappear by PND14, type 1 fibers and fibro-adipogenic progenitors recede while 2B fibers expand, ultimately constituting over three-quarters of the nuclei in gastrocnemius by 2 months (Fig. S5). Given that the majority of dynamic cCREs are cell-type specific (Fig. 4g) and the predominant cell-type-specific cCREs are found in myonuclei (Fig. 4d), we then focused on TF binding in myonuclear subtypes. We performed motif enrichment analysis using ArchR ^77^ in myonuclei-specific cCREs, classified by their accessibility in muscle fibers and satellite cells, to identify potential regulators which were then matched to TFs featured in our topic modeling (Methods, Fig. S12). Notably, some TFs exhibited concordant motif activity patterns and topic weight. The *Pax7* motif is enriched in satellite-specific cCREs (Fig. 4h) and also included in the satellite-associated topics (Fig. 4i). This is fully consistent with expectations from known biology. Alternatively, *Myog* binding was detected and the TF found highly weighted in one major satellite topic (GC15, 44% participation in satellites), whereas it is not detected in the minor satellite topic (GC8, 12% participation) (Fig. 4h,i, S12). The more dominant topic potentially reflects satellite cells actively undergoing postnatal myogenic differentiation, while the minor topic may signify the self-renewing pool of satellite cells that actively inhibit the expression of myogenin and related MRFs ^44,45^. Previous studies have found interactions between *Tcf12* and *Mef2c* and MRFs such as *Myod1* in skeletal muscle implicated in skeletal muscle formation ^78–82^. While *Tcf12* was weighted in nearly all myonuclear topics, its homodimer motif enrichment showed highest activity in satellite cells, in which previous studies have shown it to be a crucial regulator of chromatin remodeling ^78^, whereas the heterodimer motif is weakly enriched in Type 1 and Type 2 myonuclei. Similarly, *Mef2c* is found in all non-satellite topics but its motif-inferred activity is only in type 1 myonuclei. *Mef2c* has indeed been linked to type 1 specification by responding to calcium-dependent signaling pathways that alter Mef2 protein post-translationally where it acts to promote the transition between fast glycolytic fibers to slow oxidative fibers ^79–82^. In both cases, integrating accessibility and motif enrichment suggests how known post-transcriptional controls of specific TFs can parse muscle RNA topics, and from this we can make testable predictions about cCREs that are likely involved.

### Comparison of sex-specific regulatory activity in the adrenal zona fasciculata

We then turned to sex-specific cCREs that are also cell-type-specific in adrenal gland (Fig. 4j). Unsurprisingly, female cCREs overlap those attributed to the X-zone and zona fasciculata (Fig. 4k), as well as adipocytes. In males, a faint signature is seen in the nuclei annotated as male ZF. We focused motif enrichment on the X-zone, male ZF, and non-sex-specific ZF to investigate binding activity of key TFs from differential expression analysis and topics modeling. *Runx2*, upregulated in female compared to male ZF, has distinct binding activity in X-zone-specific cCREs (Fig. 4l). It is also a top-weighted gene in the X-zone topic AD6 (Fig. 4m). Despite a previous study in *Runx2* knockout mice suggesting no direct contribution to sex determination ^83^, it may regulate genes involved in steroid metabolism, as evidenced in mouse osteoprogenitor cells ^84^. Furthermore, estrogen receptor alpha has been observed to colocalize with Runx2 in breast cancer and osteoblasts, although their expression is inversely related ^85^. In contrast to *Runx2*, *Thrb* is also differentially upregulated in female ZF but is weighted similarly in X-zone topic AD6 and male ZF topic AD12 with binding activity solely in the male ZF (Fig. 4l, m). Likewise, the androgen receptor gene *Ar* is highly weighted in both the X-zone topic AD6 as well as male ZF topic AD12, but only active in male ZF (Fig. 4l, m). *Ar* is expressed in both male and female sex-specific regions, although more so in the X-zone compared to the male-specific ZF (Fig. S1). Recent studies have identified androgen signaling via the androgen receptor as a requirement for X-zone regression during puberty in male mice ^86^, while *Ar* signaling is not essential for regression in female mice ^87^. Our results suggest androgen signaling in male ZF may be mediated by lower levels of *Ar* compared to female ZF, perhaps due to co-activator expression, accessible chromatin at target gene promoters, or involvement of factors from other tissues, such as the hypothalamic-pituitary-gonadal axis. More broadly, the sexual dimorphic binding activity of transcription factors that are similarly expressed in these homologous cells highlights the fundamental limitations of studying gene regulation using RNA expression alone when ignoring sex as a biological variable.

## Discussion

The ENCODE4 mouse single-nucleus dataset stands out from other genomic catalogs by offering a comprehensive map of postnatal development across diverse tissues, spanning from just after birth to late adulthood in both sexes. This inclusivity allowed us to analyze sexual dimorphism across time, as in the example of sex-specific adrenal cortex populations during puberty. The dataset facilitated comparison of maturation rates across tissues, revealing significant differences. For instance, the most significant changes in the adrenal gland occur between 2 months and 18-20 months as sex-specific cortical layers regress, while the largest changes in gastrocnemius occur from postnatal day 4 to postnatal day 10 as myofibers mature. A time course at this resolution enables investigations into large-scale dynamics as well as the maintenance of adult stem cell pools like OPCs, NPCs, and satellite cells. Additionally, integration of snRNA-seq data between Parse and 10x barcoding platforms underscores the complementary information captured by each technology. In summary, this dataset presents a unique opportunity to explore postnatal development throughout the entire mouse body at unprecedented single-cell resolution, offering insights from various biological and technical perspectives.

All experiments were conducted in a B6/CAST hybrid genotype, facilitating future exploration of the genetic basis of complex molecular traits. B6J (*M. m. domesticus*), which is the most commonly used laboratory mouse and the first murine genome published, diverged from CAST (*M. m. castaneus*) approximately one million years ago ^88,89^. As a wild-derived strain, CAST harbors 17.6 million single-nucleotide polymorphisms relative to the B6J reference genome ^90^ and it exhibits phenotypic differences in behavior and hearing ability ^91^. These strains represent broader genetic diversity, resembling natural populations, and are two of the founders of the Collaborative Cross ^92^. An open question is whether any of the cell states described here would be specific to the F1. Examining gene expression differences in both B6 and CAST parents with our results in the offspring could allow us to determine the impact of a particular allele as acting in *cis* or *trans* ^93^. Besides allele-specific gene expression, we could also compare traits such as proportions and dynamics of cell types, as well as participation in the regulatory topics described here, streamlining the identification and analysis of cell types and states.

We applied Topyfic to integrated combinatorial barcoding and multiome snRNA-seq datasets, focusing on a curated vocabulary of 2,701 regulatory genes. We recovered 82 regulatory topics associated with 46 distinct cell types and states. Our dataset shows the strength of topic modeling for capturing cell-level changes within clusters, as described for differentiating cell types such as oligodendrocytes and DG neurons. Our regulatory topics allowed us to study the biotypes of genes that change from one topic to another as well as compare topics learned independently in separate tissues. Our results indicated an enrichment of transcription factor (TF) and microRNA gene biotypes in cell-type-specific topics, while cycling topics are predominantly influenced by chromatin regulators. Although most studies of polyadenylated RNA ignore the impact of microRNAs, a significant fraction of microRNAs are intragenic, most of which are found within introns of protein-coding genes ^94,95^. MicroRNAs can be transcribed by RNA polymerase II together with their host genes ^96^. One possibility is that microRNAs embedded in the introns of known cell type markers may play a role in the regulation of expression levels within that cell type.

Additionally, our analysis identified correlated regulatory topics across tissues for shared cell types, such as endothelial cells, while some immune cell types retained a tissue-specific signature, particularly in trunk organs compared to brain microglia. We further classified ENCODE v4 cCREs based on accessibility in our cell types, revealing that nearly half of the identified cCREs exhibit cell type specificity. Lastly, we explored motif enrichment patterns of TFs within topics in cell type- and state-specific regulatory elements.

The behavior of rLDA topics aligns with our expectations about genuine cellular programs: they are predominantly cell type- and state-specific, often co-expressed, reproducible across tissues, and can be defined using regulatory genes alone, especially TFs. Focusing on regulatory genes offers direct insight into cellular programs by ensuring the inclusion of TFs in each topic rather than putting higher weights on downstream targets, many of which encode structural proteins or have no known function. It is also likely that there will be interesting differences in specific topics in different mouse strains as well as possibly altogether new topics for cell states not present in our F1 mice. It is crucial to note that a TF’s presence in a topic does not automatically imply active involvement in regulatory programs, and further verification may require follow-up experiments and integration with chromatin accessibility or DNA binding data. By leveraging corresponding chromatin accessibility data, we identified cases where a top-weighted TF exhibits enriched binding in a cell type associated with its topic, as well as instances where topic TFs are active in different cell types or states. Our results demonstrate the successful identification and interpretation of cellular programs using topic modeling across multiple tissues and barcoding platforms, establishing a foundation of non-exclusive transcriptional programs operating across postnatal development. We also showed they can be linked to downstream *cis*-regulatory targets.

## Data and code availability

- **Data availability**: ENCODE carts of all data used are listed in Table S1.
- **Data processing/figure generation code**: https://github.com/erebboah/enc4_mouse_paper/

## Acknowledgements

We thank the Caltech Jacobs Genetics and Genomics Laboratory for sequencing the bulk mRNA-seq libraries for Illumina sequencing and the UCI GHTF for PacBio sequencing. A.M. and B.J.W. were supported by UM1HG009443. B.J.W. was also supported by the Caltech Beckman Institute BIFGRC. J.J and I.Y. (ENCODE DCC) were supported by U24HG009397. M.P.S. was supported by 1UM1HG009442.

## Author contributions

E.R. performed Parse Biosciences snRNA-seq experiments, microRNA-seq experiments, data processing, data analysis, generated figures, wrote and edited the manuscript with significant input from B.J.W. and A.M. N.R. performed data processing, data analysis, generated figures, wrote and edited the manuscript. B.A.W. processed RNA samples, performed bulk mRNA-seq experiments, wrote and edited the manuscript. A.W., M.S., and X.Y. performed 10x Multiome experiments. H.Y.L. performed Parse Biosciences snRNA-seq experiments and sequenced snRNA-seq and microRNA-seq libraries. L.A.D. and L.R. bred mice, dissected tissues, and shipped samples to Caltech. S.M. performed data processing and F.R. edited the manuscript. F.R., D.T., J.J., and I.Y. contributed to ENCODE uniform processing pipeline development for Parse Biosciences snRNA-seq data. All authors read and approved the final manuscript.

## Methods

### Mice and tissue collection

All animals were treated and housed in accordance with the Guide for Care and Use of Laboratory Animals. Approval for all experimental procedures was granted by Caltech’s Institutional Animal Care and Use Committee (IACUC), aligning with both institutional and national guidelines. Samples were obtained from animals covered under the approved IACUC protocol #IA21-1647, “Single-cell transcriptome studies from multiple mouse tissues”. Tissues at postnatal day (PND) 4, PND 10, PND 14, PND 25, PND 36, 2 months, and 18-20 months from C57BL6/J (RRID:IMSR_JAX:000664) × CAST/EiJ (RRID:IMSR_JAX:000928) F1 hybrid mice were obtained from Jackson Laboratories (JAX). Adrenal gland and gastrocnemius tissues were pooled from 3 individuals for PND 4 and PND 10 timepoints. Hippocampus tissues were pooled from 3 individuals for PND 10 and PND 14 timepoints. Tissues were flash-frozen in liquid nitrogen and delivered to Caltech on dry ice, where they were stored at −80°C until RNA extraction.

### Isolation of RNA for bulk assays

For bulk RNA-seq, total RNA was extracted from flash-frozen tissues at Caltech using the Norgen Animal Tissue RNA Purification Kit (Norgen Biotek cat. #25700). The tissue was lysed using Buffer RL and proteins were digested with proteinase K. Genomic DNA was removed with DNaseI treatment on the columns. The purified total RNA includes includes large mRNAs, lncRNAs, and small RNAs. The Qubit dsDNA HS Assay Kit (Thermo cat. #Q32854) was used to assess RNA concentration and RIN values were determined using the Bioanalyzer Pico RNA kit (Agilent cat. #5067-1513), with average RIN scores of 8.2 for the adrenal gland, 9.1 for the hippocampus, 9.3 for the cortex, 9.0 for the heart, and 9.3 for gastrocnemius tissues.

### Bulk RNA-seq from mouse tissues

Each cDNA library was built from 300 ng total RNA with ERCC spike-ins (Thermo cat. #4456740) using the NEBNext Ultra II Directional RNA Library Prep Kit for Illumina (NEB cat. #E7760), specifically the protocol for use with NEBNext Poly(A) mRNA Magnetic Isolation Module (NEB cat. #E7490). Ribosomal RNA was depleted from total input RNA using the NEBNext rRNA Depletion Kit (NEB cat. #E6310). First and second strand synthesis, cDNA end prep, adapter ligation, and finally PCR amplification resulted in the final libraries. The libraries were quantified using the Qubit dsDNA HS Assay Kit (Thermo cat. #Q32854) and sequenced on an Illumina HiSeq 2500 as 100 bp single-end reads to 50 M raw read depth. For submission to the ENCODE portal, libraries needed at least 30 M aligned reads and a Spearman replicate correlation *>*0.9.

### Purification of nuclei for Split-seq

For Parse Split-seq experiments performed at UCI, nuclei were isolated from the 5 core tissues (adrenal gland, left cerebral cortex, hippocampus, heart, and gastrocnemius) for all 7 timepoints (PND 4, PND 10, PND 14, PND 25, PND 36, 2 months, and 18-20 months). Flash-frozen tissues shipped from Caltech were transferred to a chilled gentleMACS C Tube (Miltenyi Biotec cat. #130-093-237) with 2 mL Nuclei Extraction Buffer (Miltenyi Biotec cat. #130-128-024) supplemented with 0.2 U/uL RNase Inhibitor (NEB cat. #M0314L) on ice. Nuclei were dissociated from whole tissues using a gentleMACS Octo Dissociator (Miltenyi Biotec cat. #130-095-937). Suspensions were filtered through a 70 um strainer then a 30 um strainer (Miltenyi Biotec cat. #130-110-916 and #130-098-458, respectively). Nuclei were resuspended in cold PBS + 7.5% BSA (Life Technologies cat. #15260037) and 0.2 U/ul RNase inhibitor for manual counting using a hemocytometer and DAPI stain (Thermo cat. #R37606). For gastrocnemius tissue, debris was removed from nuclei suspensions with Debris Removal Solution (Miltenyi Biotec cat. #130-109-398). Nuclei were mixed with Debris Removal Solution and layered on top of PBS, then centrifuged at 4°C, 3000 x g for 10 minutes with full acceleration and no brake. Nuclei bands were separated from debris layers and concentrations were determined using a hemocytometer. For Parse Split-seq, 1-4 million nuclei per sample were fixed using Parse Biosciences’ Nuclei Fixation Kit v1 (Parse Biosciences cat. #WN100), following the manufacturer’s protocol. Briefly, nuclei were incubated in fixation solution for 10 minutes on ice, followed by permeabilization for 3 minutes on ice. The reaction was quenched, then nuclei were centrifuged and resuspended in 300 uL Nuclei Buffer (Parse Biosciences cat. #WN101) for a final count. DMSO (Parse Biosciences cat. #WN105) was added before freezing fixed nuclei at −80°C.

### Parse Split-seq experiments

Nuclei were barcoded using Parse Biosciences’ Evercode WT Kit v1 (cat. #EC-W01030), following the manufacturer’s protocol. Briefly, fixed nuclei were thawed and added to the Round 1 reverse transcription barcoding plate at 15,000 nuclei per well across 48 wells. Individual samples from each tissue were distributed in sample barcoding plates with at least 1 well per individual. Within the fixed nuclei, RNA was reverse transcribed using oligodT and random hexamer primers and the first barcode was annealed. After RT, nuclei were pooled and distributed in 96 wells of the Round 2 ligation barcoding plate for in situ barcode ligation. After Round 2, nuclei were pooled and redistributed into 96 wells of the Round 3 ligation barcoding plate for barcode 3 and Illumina adapter ligation. Finally, nuclei were counted using a hemocytometer and distributed into 6 subpools for adrenal, 6 subpools for cortex, 5 subpools for hippocampus, 4 subpools for heart, and 5 subpools for gastrocnemius, each containing 12,000 nuclei, with 2 additional subpools of 15,000 nuclei for gastrocnemius. Nuclei from each tissue were also distributed into 1-2 small subpools of 1,000-2,000 nuclei each, for a target of around 75,000 nuclei per tissue (*>*500 UMI). The nuclei in each subpool were lysed and the barcoded cDNA underwent template switching and amplification. The cDNA was cleaned using AMPure XP beads (Beckman Coulter cat. #A63881) and quality checked using the Qubit dsDNA HS Assay Kit (Thermo cat. #Q32854) and a Bioanalyzer 2100 (Agilent cat. # G2939A) High Sensitivity DNA Kit (Agilent cat. #5067-4626) before proceeding to Illumina library preparation with 100 ng of full-length cDNA per subpool. Subpool cDNA was fragmented and Illumina P5/P7 adapters were ligated during the final amplification, followed by size selection and quality check with the Bioanalyzer and Qubit. Libraries with 5% PhiX spike-in were sequenced on an Illumina NextSeq 2000 sequencer with P3 200 cycles kits (Illumina cat. #20040560) as paired-end, single-index reads (115/86/6/0) to an average depth of 181 M reads per 12,000-15,000-nucleus library and an average depth of 134 M reads per 1,000-2,000-nucleus library.

### Purification of nuclei for 10x Multiome

For 10x Multiome experiments performed at Stanford University, nuclei were isolated from 5 core tissues for PND 14 and 2 month timepoints. Flash-frozen tissues were dissociated in a Douce homogenizer with 1 mL homogenization buffer: 0.26 M sucrose (Sigma cat. #S7903-250G), 0.03 M KCl (Thermo cat. #AM9640G), 0.01 M MgCl2 (Thermo cat. #AM9530G), and 0.02 M Tricine-KOH pH 7.8 (Sigma cat. #T0377), supplemented with 0.6 U/uL RNase Inhibitor (Thermo cat. #EO0384). Suspensions were filtered through a 40 um strainer (Fisher Scientific cat. #22363547) and debris was removed using an iodixanol gradient. Iodixanol solution was diluted from 60% iodixanol (Sigma cat. #D1556-250ML) with dilution buffer consisting of 0.15 M KCl, 0.03 M MgCl2, and 0.12 Tricine-KOH pH 7.8. Nuclei were mixed 1:1 with 50% iodixanol solution, then 30% iodixanol solution was layered underneath the 25% mixture, and 40% iodixanol solution was layered at the bottom. Nuclei were centrifuged at 4°C, 3000 x g for 20 minutes with full acceleration and no brake and the nuclei band was separated from the debris layer. Concentrations of the final suspensions were determined using a hemocytometer. Nuclei were immediately processed following the Chromium Next GEM Single Cell Multiome ATAC + Gene Expression User Guide (CG000338).

### 10x Multiome experiments

Gene expression and chromatin accessibility were profiled simultaneously in the same nuclei using the Chromium Next GEM Single Cell Multiome ATAC + Gene Expression kit (10x Genomics cat. #1000283) following the manufacturer’s protocol. Briefly, around 16,000 nuclei were loaded per well in the microfluidic chip and partitioned into gel beads-in-emulsions (GEMs) for a target recovery of 5,000-10,000 nuclei per sample (around 80,000 nuclei per tissue). During incubation, transposase cleaved open regions of DNA and added GEM-specific adapter sequences to the fragments. After transposition, the nuclei lysates were reverse transcribed using oligodT primers, which also adds GEM-specific barcodes and UMIs to the resulting cDNA. The GEMs were then broken and the transposed DNA and barcoded cDNA underwent pre-amplification PCR to produce the input material for parallel snATAC-seq and snRNA-seq library building. For snATAC-seq, Illumina P5/P7 adapters were added during sample index PCR and the final libraries were cleaned using SPRIselect beads (Beckman Coulter cat. #B23318). For snRNA-seq, the barcoded cDNA underwent template switching and amplification, and was then fragmented and size-selected using SPRIselect beads. Illumina P5/P7 adapters were added during sample index PCR and the final snRNA-seq libraries were cleaned using SPRIselect beads. The snATAC-seq libraries were sequenced on an Illumina NovaSeq 6000 sequencer as paired-end, dual-indexed reads (50/50/8/24) to an average depth of 180 M reads per library. The snRNA-seq libraries were sequenced on an Illumina NovaSeq 6000 sequencer as paired-end, dual-indexed reads (28/90/10/10) to an average depth of 194 M reads per library.

### Demultiplexing Parse Biosciences snRNA-seq data

Due to the combinatorial barcoding approach, raw fastqs from Parse snRNA-seq libraries contain all samples included in the experiment. In order to provide sample-level fastqs to the ENCODE portal, Parse Biosciences’ split-pipe software v0.7.6p and custom code were used to assign reads to samples. Briefly, split-pipe v0.7.6p was used to generate an annotated fastq with read names containing cell barcodes (process/single_cells_barcoded_- head.fastq.gz) as well as a cell metadata file (all-well/DGE_unfiltered/cell_metadata.csv) mapping barcode to sample for each pair of subpool fastqs associated with an experiment. A custom python script calls seqtk v. 1.3-r106 (https://github.com/lh3/seqtk) to extract reads from the original fastqs and output them as sample-level fastq files.

### Read mapping and quantification

All data quantifications were downloaded from ENCODE portal using carts, organizing the data based on assay and/or tissue (refer to Table S1 for links to carts).

Bulk and single-nucleus RNA-seq data were processed through ENCODE uniform processing pipelines using the mm10 genome with Gencode vM21 annotations. For bulk RNA-seq, the data were aligned using STAR v. 2.5.1b ^97^ and quantified using RSEM, which provides FPKM, TPM, and raw counts (https://www.encodeproject.org/pipelines/ENCPL862USL/).

The snRNA-seq data were aligned using STARSolo v. 2.7.10a ^98^ with GeneFull_Ex50pAS settings to generate UMI count matrices (https://www.encodeproject.org/pipelines/ENCPL257SYI/), similar to the intronic count option in 10x’s Cell Ranger.

Single-nucleus ATAC-seq data were processed using the standard ENCODE snATAC-seq pipeline with the mm10 genome to generate fragment files which were used as input to downstream analyses (https://www.encodeproject.org/pipelines/ENCPL952JRQ/).

### Bulk RNA-seq analysis

Normalized bulk RNA-seq quantifications were concatenated across all samples using the TPM column from the ENCODE pipeline. In each tissue, the number of regulatory genes in each category were counted if they were expressed at *>*1 TPM in at least 1 bulk sample.

### QC and filtering of single-nucleus data

Analyses were performed on a per-tissue basis and all input files were downloaded from the ENCODE portal. The snRNA-seq tar files contain sparse matrices with corresponding gene and barcode CSV files. The corresponding snATAC-seq tar files for 10x Multiome contain compressed TSV fragments and indices. For Parse Split-seq, the number of datasets varies depending on the number of subpools set aside per tissue.

To perform the integrated snRNA-seq analysis, 42 Parse Split-seq datasets and 8 10x Multiome datasets for adrenal gland, 32 Parse Split-seq datasets and 8 10x Multiome datasets for cortex, 34 Parse Split-seq datasets and 8 10x Multiome datasets for hippocampus, 28 Parse Split-seq datasets and 8 10x Multiome datasets for heart, and 56 Parse Split-seq datasets and 8 10x Multiome datasets for gastrocnemius were downloaded from the ENCODE portal (Table S3). Genes were filtered for protein coding, lncRNAs, pseudogenes, and microRNAs. Ambient RNA was filtered from droplet-based 10x data using Cellbender v. 0.2.2^99^. Doublet detection was performed on nuclei with *>* 500 UMIs detected per nucleus using Scrublet v. 0.2.3^100^.

Data were filtered differently for the “standard” Parse Split-seq libraries (12,000-15,000-nucleus subpools), small Parse Split-seq libraries (1,000-2,000-nucleus subpools), and 10x Multiome nuclei (5,000-nucleus libraries). The Parse Split-seq nuclei belonging to the 12-15,000-nucleus subpools were filtered by *>* 500 and *<* 30,000 UMIs per nucleus, *>* 500 genes expressed, *<* 0.2 doublet score, and *<* 0.5 percent mitochondrial gene expression for adrenal gland, cortex, and hippocampus, and the 1-2,000-nucleus subpools by *>* 1000 and *<* 50,000 UMIs. For heart, the filters were relaxed slightly to *<* 0.25 doublet score and *<* 1 percent mitochondrial gene expression and further relaxed for gastrocnemius to *<* 5 percent mitochondrial gene expression. The 10x Multiome nuclei were filtered slightly differently: *>* 500 and *<* 30,000 UMIs, *>* 300 genes, *<* 0.25 doublet score, and *<* 5 percent mitochondrial gene expression for cortex, hippocampus, and gastrocnemius, and *>* 1000 UMIs, *<* 0.2 doublet score, and *<* 0.5 percent mitochondrial gene expression for adrenal gland and heart. In addition, 10x Multiome nuclei were also filtered by *>* 1000 unique nuclear fragments, TSS enrichment *>* 4, and *<* 1 ArchR doublet score in the corresponding snATAC-seq data. After initial processing of snATAC-seq data (described below), barcode sequences from snRNA-seq and snATAC-seq multiome nuclei were matched and nuclei failing snATAC-seq QC were excluded from downstream snRNA-seq analysis. All filtering parameters per library can be found in Table S2.

### Preprocessing 10x snATAC-seq data

ArchR Arrow files were generated for each tissue using the ENCODE processed fragments files from 8 experiments with a minimum TSS enrichment of 4, minimum 1,000 unique fragments per cell, and excluding reads from mitochondrial DNA in downstream analysis ^77^. Doublets were scored and filtered using ArchR’s “addDoubletScores” and “filterDoublets” functions with an enrichment threshold of 1^77^. ArchR projects for each tissue were saved and barcode sequences were translated into their snRNA-seq counterpart and saved as csv files. After snRNA-seq filtering, nuclei failing snRNA QC were dropped from the ArchR project using “subsetArchRProject”.

### Integration of Parse and 10x snRNA-seq data

After filtering the 3 Seurat objects per tissue (standard Parse, small Parse, and 10x Multiome), each was normalized using the function “SCTransform” in Seurat v. 4.1.1^101^, with number of genes expressed per nucleus and percent mitochondrial gene expression regressed out. Anchors for integration across the 3 objects were calculated using “SelectIntegrationFeatures” with 3,000 genes, “PrepSCTIntegration”, and “FindIntegrationAnchors” in Seurat, with the standard Parse dataset serving as the reference due to inclusion of all 7 timepoints. After integrating data (“IntegrateData”), principal component analysis was performed on the integrated assay by the “RunPCA” function with 50 principal components, with the UMAP (“RunUMAP”) calculated from the first 30 components. Clustering was performed with the Louvain clustering algorithm (“FindClusters”) with resolution 0.8, with sub-clustering performed as necessary on specific clusters in gastrocnemius and hippocampus due to expression of known marker genes (Fig. S3, S5).

### Integrated cell type annotation

When available, reference datasets were used to transfer annotations using “FindTransferAnchors” in Seurat v. 4.1.1^101^. For both cortex and hippocampus, a downsampled version of the 1M whole cortex and hippocampus 10x atlas from 8 week old mice available on the Allen data portal ^10^ was used to transfer subtype-level annotations. Downsampling was performed per “cell_type_alias_label” group, with 1,000 nuclei taken per cell type (or all nuclei, if *<* 1,000 were available) for a total of 250,734 nuclei used for label transfer. For the heart dataset, both a human heart cell atlas ^23^ (486,134 nuclei) and a dataset of 8-14 week old stressed mouse ventricles ^24^ (29,615 nuclei) were used separately for label transfer. For gastrocnemius, label transfer was performed using P10, P21, and 5-month mouse tibialis anterior datasets ^6^ (28,047 total nuclei). In addition to label transfer, curated marker genes were used to refine predictions (Fig. S1, S2, S3, S4, S5, Table S3). In lieu of a reference dataset in the case of adrenal gland, marker genes alone were used to annotate celltypes per cluster. Each cluster was annotated at the finest possible resolution in a grouping titled “subtypes” (in all figures, metadata, and data objects). This resolution includes dynamic cell states such as OPCs, early DG, the sex-specific populations in the adrenal cortex, and layer-specific neuronal subtypes in cerebral cortex. Depending on the downstream analysis, subtypes and states were grouped into a coarser resolution titled “celltypes”. For example, transient sex-specific populations in the adrenal cortex are collapsed along with zona fasciculata, and cerebral cortex layers are all annotated as glutamatergic neurons.

### Transferring cell type annotations to corresponding snATAC-seq

Cell type annotations were added to each ArchR project using the per-cell metadata extracted from Seurat objects. Barcode sequences were matched between assays and annotations carried over from snRNA-seq analysis with no modifications.

### Differential gene expression analysis of pseudobulk snRNA-seq

The raw, unnormalized counts were extracted from the annotated Seurat object for subtypes of interest and summed across all nuclei in each individual mouse for a sample-level pseudobulk counts matrix across all expressed genes. Using pydeseq2^102^, defined groups such as sex were compared within subtypes. Results were filtered by an absolute log fold change *>*1 and adjusted p-value *<* 0.01.

### Pseudotime ordering of dynamic cell states in hippocampus

Cell types of interest were subset from the tissue-level Seurat object for pseudotime ordering using Monocle 3^50,103–106^. The root cells were chosen for “order_cells” according to the known stage of the cells. The oligodendrocytes and OPCs were subset from the hippocampus dataset, with root cells corresponding to the OPCs. For ordering of the DG cells, root cells correspond to the cells from early timepoints. Pseudotime values for the ordered cells were incorporated into their metadata for downstream analysis.

### Calculating single-nucleus regulatory topics using Topyfic

The raw, unnormalized counts were extracted from each filtered Seurat object per tissue and barcoding technology (Parse and 10x). Genes were filtered to 2,701 regulatory genes ^19^ determined by microRNA-host gene correlations, annotated transcription factors, and genes annotated with the following Gene Ontology (GO) terms: 0004402 (histone acetyltransferase activity), 0004407 (histone deacetylase activity), 0042054 (histone methyltransferase activity), 0032452 (histone demethylase activity), 0016592 (mediator complex), 0006352 (DNA-templated transcription, initiation), 0003682 (chromatin binding), 0006325 (chromatin organization), 0030527 (structural constituent of chromatin), and 0140110 (transcription regulator activity). MicroRNA host genes were included if they are annotated as a host gene (e.g. *Mir133a-1hg*, *Mir124a-1hg*) and/or their Spearman correlation with expression of the mature microRNA was *≥* 0.3^19^.

Depth normalization was performed on each raw counts matrix by tissue (x 5) and technology (Parse and 10x; 10 total matrices) by a round of proportional fitting followed by log transformation, then another round of additional proportional fitting ^107^. An anndata object was constructed from the normalized matrix, 2,701 regulatory genes, and per-cell metadata including subtype and celltype annotations.

Topyfic was run with a range of *k* values for each tissue and technology using 100 runs of LDA with batch_size of 128 and 5 minimum iterations ^19^. The best k per tissue and technology was determined by comparing *k* to the number of resulting topics, *n*. The closest *k* to the resulting *n* value was chosen: *k* = 15 for Parse and 13 for 10x adrenal, 14 for Parse and 13 for 10x cortex, 13 for Parse and 21 for 10x hippocampus, 11 for Parse and 13 for 10x heart, and 12 for Parse and 8 for 10x gastrocnemius. Harmony ^55^ was used to combine the best models learned separately from each technology to a unified set of topics, filtering out topics with participation in less than 1% of nuclei in the smaller of the two datasets. Downstream analysis such as comparisons between topics was facilitated by analysis of the gene weights in each topic (Tables S4-S8).

### Topics analysis

Harmonized snRNA-seq topics in each tissue were characterized by analysis of topic-trait enrichment (Topyfic function “TopicTraitRelationshipHeatmap” on the analysis TopModel object), a measurement of how highly-weighted topic genes are specifically expressed in traits like celltypes, subtypes, ages, and sexes ^19^. Topics were further interpreted by cell participation across celltypes and subtypes, represented as pie charts (function “pie_structure_Chart”) and structure plots (function “structure_plot”) ^19^. Two specific topics of interest, such as immune-related topics in heart and brain, were compared using an MA plot (function “MA_plot”), and topics were compared across tissues by Pearson correlation based on gene weights ^19^.

### Characterizing ENCODE cCRE specificity with snATAC-seq

The ENCODE V4 catalog of candidate cis-regulatory elements (cCREs) for mm10 was downloaded from the ENCODE portal (https://www.encodeproject.org/files/ ENCFF167FJQ/) ^76^. All 926,843 cCREs were added to each tissue’s ArchR project by the function “addPeakSet”, then scored using “addPeakMatrix”, which counts the number of fragments per region with a maximum count of 4 to prevent large biases in the counts ^77^. The raw counts matrices were extracted (“getMatrixFromProject”), pseudobulked by integrated snRNA cluster, and normalized by RPM. RPKM was not used due to the limited distribution of cCRE lengths, between 150 and 350 bp with a mean of 269 bp and standard deviation of 64.9 bp (Fig. S11). For clarity in downstream analysis, small clusters of less than 100 multiome nuclei were removed (such as a cluster corresponding to 16 hepatocytes detected in adrenal gland, most likely a dissection artifact). Each cCRE was classified as accessible in a celltype if it scored *≥* 5 RPM in at least one cluster corresponding to that celltype. Categories of “specific”, “shared”, “general”, or “global” were assigned based on the number of celltypes within and across tissues with open chromatin at each cCRE. “Specific” refers to cCREs accessible in only one celltype above the RPM threshold across all tissues. Common celltypes such as macrophages and endothelial cells were considered one celltype. “Shared” refers to cCREs accessible in more than one celltype within or across tissues. “General” refers to cCREs accessible in all major celltypes within a tissue, and “global” refers to cCREs accessible in all major celltype across all tissues. Major celltypes were defined as those whose cumulative sum makes up 90% of the cell types in the tissue; for example neurons in the brain, myonuclei in skeletal muscle, and adrenal cortical cells, followed by other major types such as glial cells, endothelial cells, and fibroblasts.

### Differential accessibility analysis of pseudobulk snATAC-seq

Pseudobulk cCRE counts matrices were generated per sample and tissue by extracting raw single-nucleus counts and summing per cCRE across all nuclei from each individual mouse. Using pydeseq2^102^, accessibility of the previously characterized cCREs accessible in pseudobulk clusters was compared between sexes and timepoints within each tissue and group, i.e. female vs. male adrenals at PND 14, female vs. male adrenals at 2 months, PND 14 vs. 2 month male adrenals, PND 14 vs. 2 month female adrenals, etc. Results were filtered by an absolute log fold change *>*2 and adjusted p-value *<* 0.01. Unique cCREs open in each group were counted and normalized by the total number of cCREs accessible in the tissue.

### Motif enrichment analysis

Motif enrichment was calculated using ArchR to analyze transcription factor activity in celltype specific cCREs. The JASPAR2024 CORE vertebrate non-redundant PFMs ^108^ were formatted as a custom RangedSummarizedExperiment, and matches with the full set of cCREs were extracted with motifmatchr ^109,110^. ArchR’s “customEnrichment” function was used to run hypergeometric-based enrichment testing on the matched motifs and a custom subset of specific cCREs as a GenomicRanges object ^77,110^. Motifs were filtered by bulk RNA-seq expression in each tissue for downstream analysis (*>*5 TPM in at least 1 sample).

## Supplementary Tables

- **Table S1: List of ENCODE portal carts for single-cell datasets grouped by tissue and assay.**
- **Table S2: Sample metadata for snRNA-seq experiments.**
- **Table S3: List of marker genes and their respective cell types in each tissue.**
- **Table S4: Gene weights in 19 adrenal gland topics.**
- **Table S5: Gene weights in 16 cerebral cortex topics.**
- **Table S6: Gene weights in 14 hippocampus topics.**
- **Table S7: Gene weights in 17 heart topics.**
- **Table S8: Gene weights in 16 gastrocnemius topics.**

## Supplementary Figures

**Figure S1.**
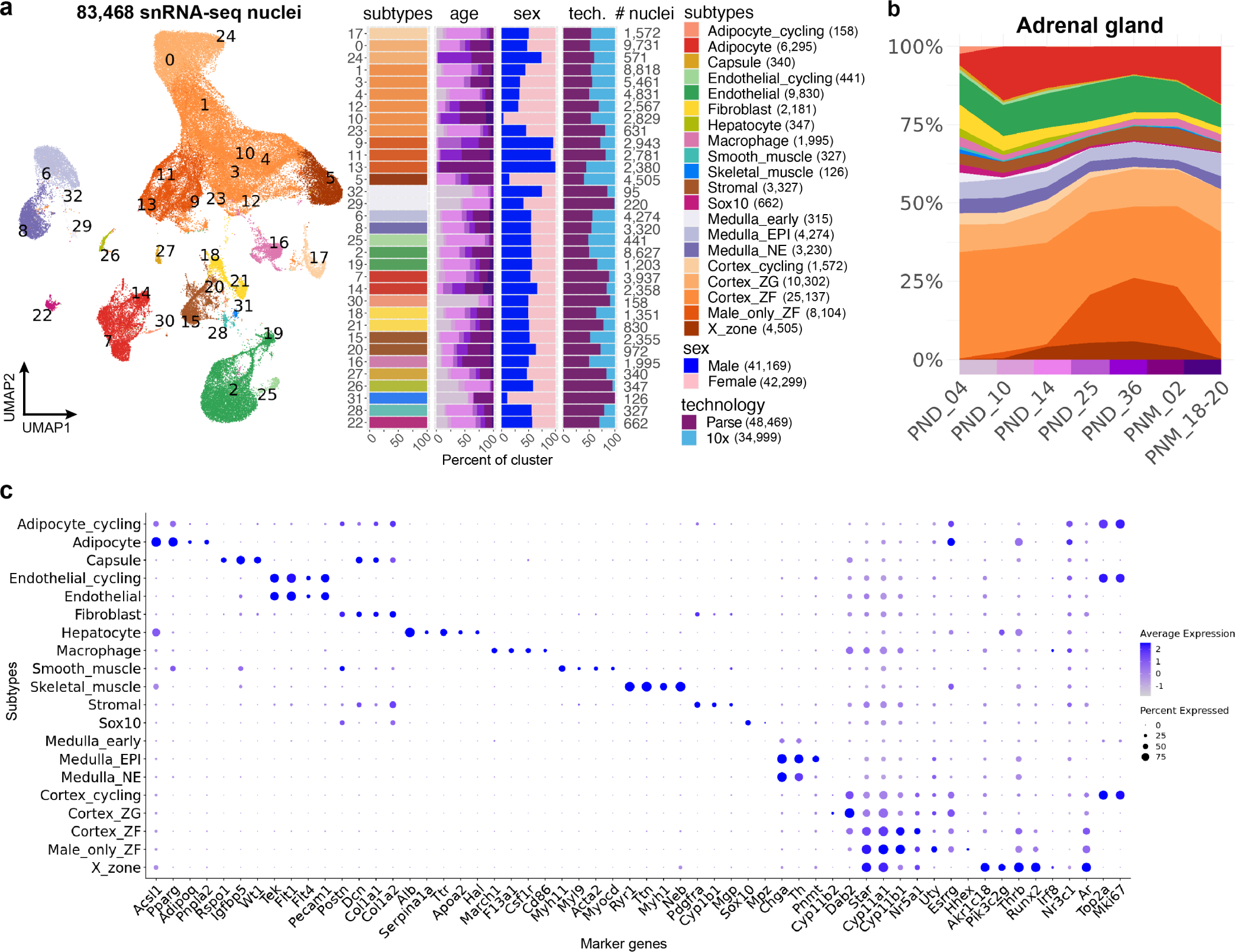
Clustering and annotation of integrated adrenal gland snRNA-seq data. **a,** UMAP representation of 83,468 adrenal gland nuclei integrated between Parse and 10x Multiome platforms and breakdown of age, sex, and technology per cluster. Numbers of nuclei per cluster are annotated to the right of the bar plots, and numbers of nuclei per annotated cell subtype are included in the legend. **b,** Dynamics of cell subtype composition across postnatal development in adrenal gland, with the same color legend as in a. For consistent sampling at each timepoint, only Parse data is shown. **c,** Expression of marker genes across subtypes in adrenal gland.

**Figure S2.**
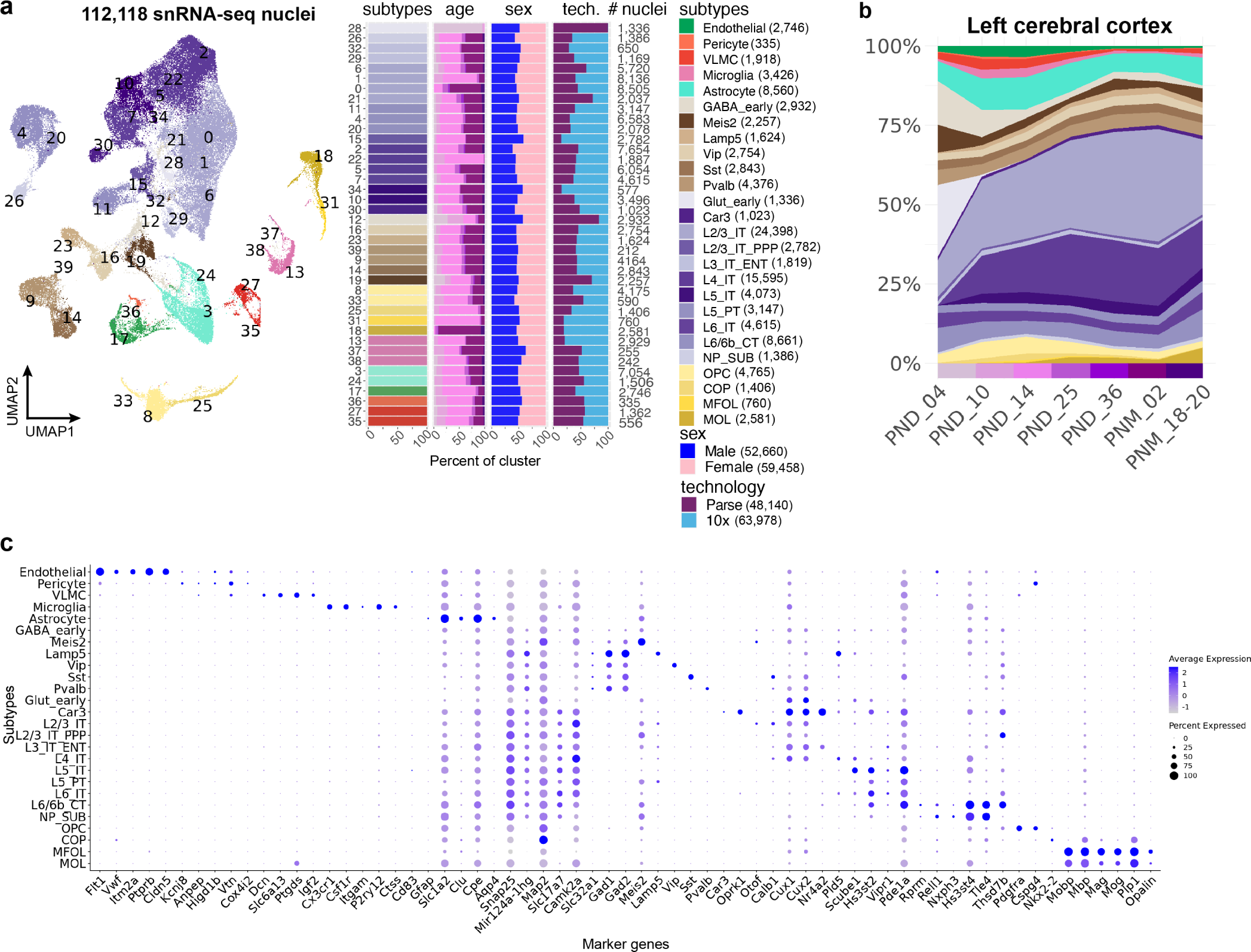
Clustering and annotation of integrated left cerebral cortex snRNA-seq data. **a,** UMAP representation of 112,118 left cerebral cortex nuclei integrated between Parse and 10x Multiome platforms and breakdown of age, sex, and technology per cluster. Numbers of nuclei per cluster are annotated to the right of the bar plots, and numbers of nuclei per annotated cell subtype are included in the legend. **b,** Dynamics of cell subtype composition across postnatal development in cortex, with the same color legend as in a. For consistent sampling at each timepoint, only Parse data is shown. **c,** Expression of marker genes across subtypes in cortex.

**Figure S3.**
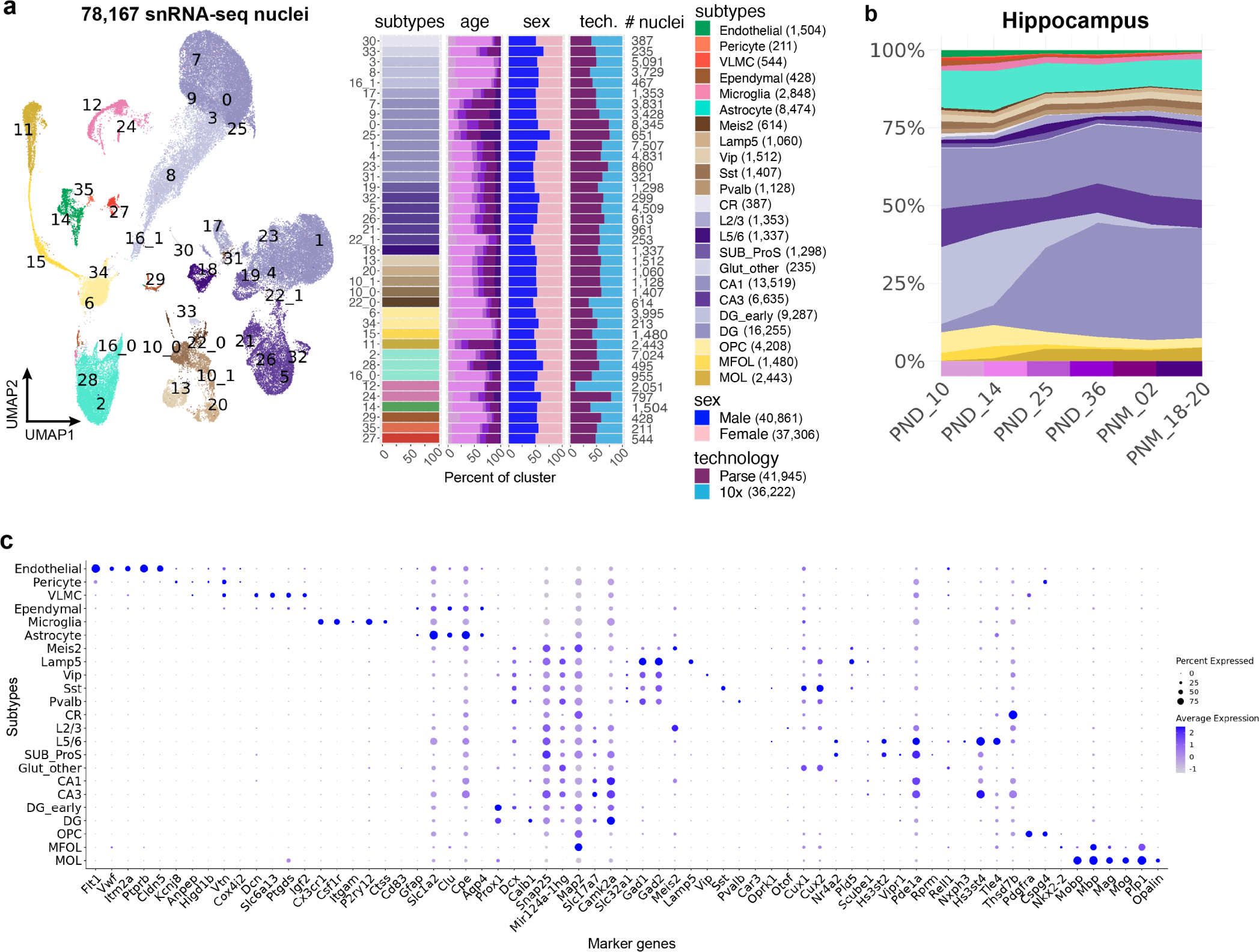
Clustering and annotation of integrated left hippocampus snRNA-seq data. **a,** UMAP representation of 78,167 left hippocampus nuclei integrated between Parse and 10x Multiome platforms and breakdown of age, sex, and technology per cluster. Numbers of nuclei per cluster are annotated to the right of the bar plots, and numbers of nuclei per annotated cell subtype are included in the legend. **b,** Dynamics of cell subtype composition across postnatal development in hippocampus, with the same color legend as in a. For consistent sampling at each timepoint, only Parse data is shown. **c,** Expression of marker genes across subtypes in hippocampus.

**Figure S4.**
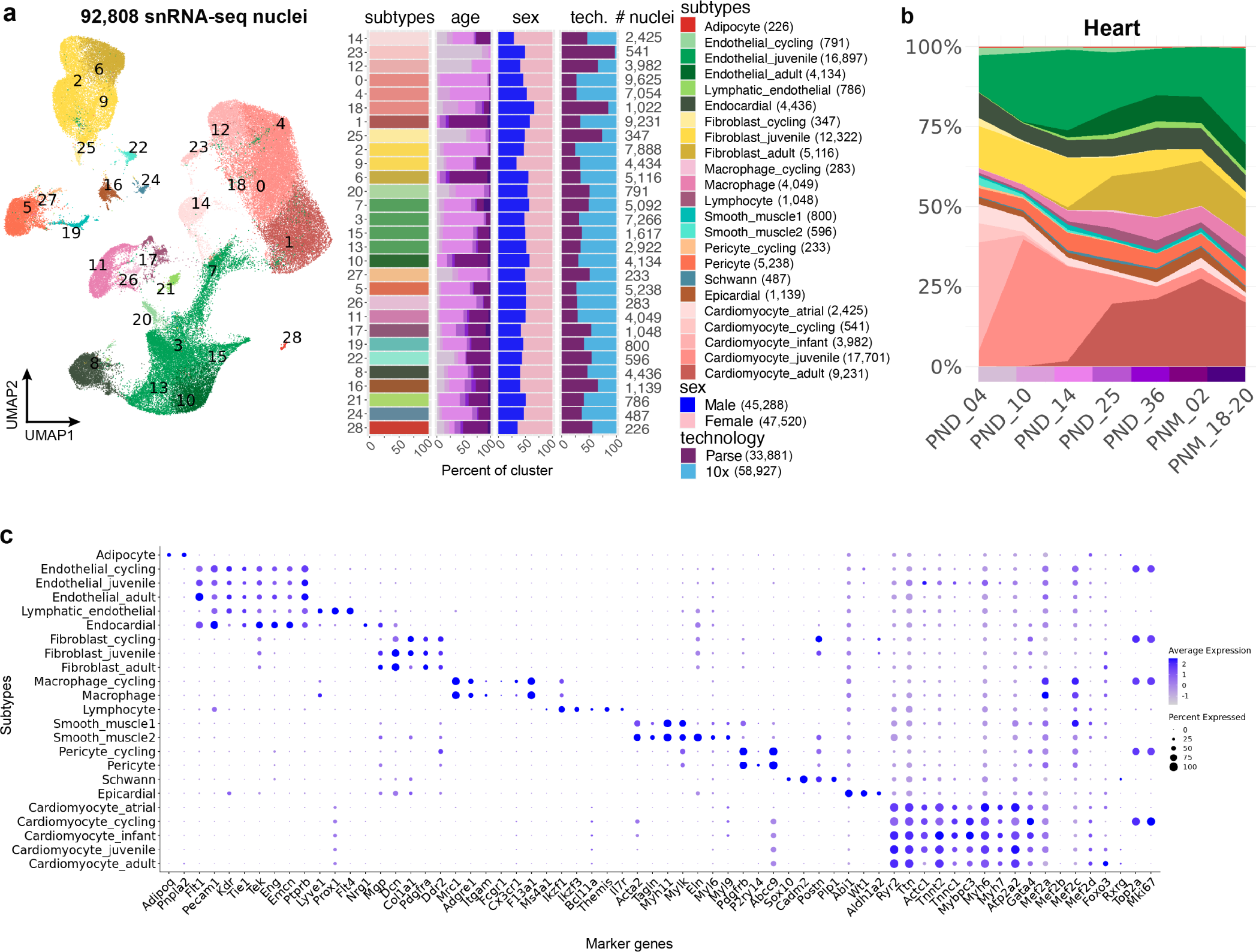
Clustering and annotation of integrated left hippocampus snRNA-seq data. **a,** UMAP representation of 78,167 heart nuclei integrated between Parse and 10x Multiome platforms and breakdown of age, sex, and technology per cluster. Numbers of nuclei per cluster are annotated to the right of the bar plots, and numbers of nuclei per annotated cell subtype are included in the legend. **b,** Dynamics of cell subtype composition across postnatal development in heart, with the same color legend as in a. For consistent sampling at each timepoint, only Parse data is shown. **c,** Expression of marker genes across subtypes in heart.

**Figure S5.**
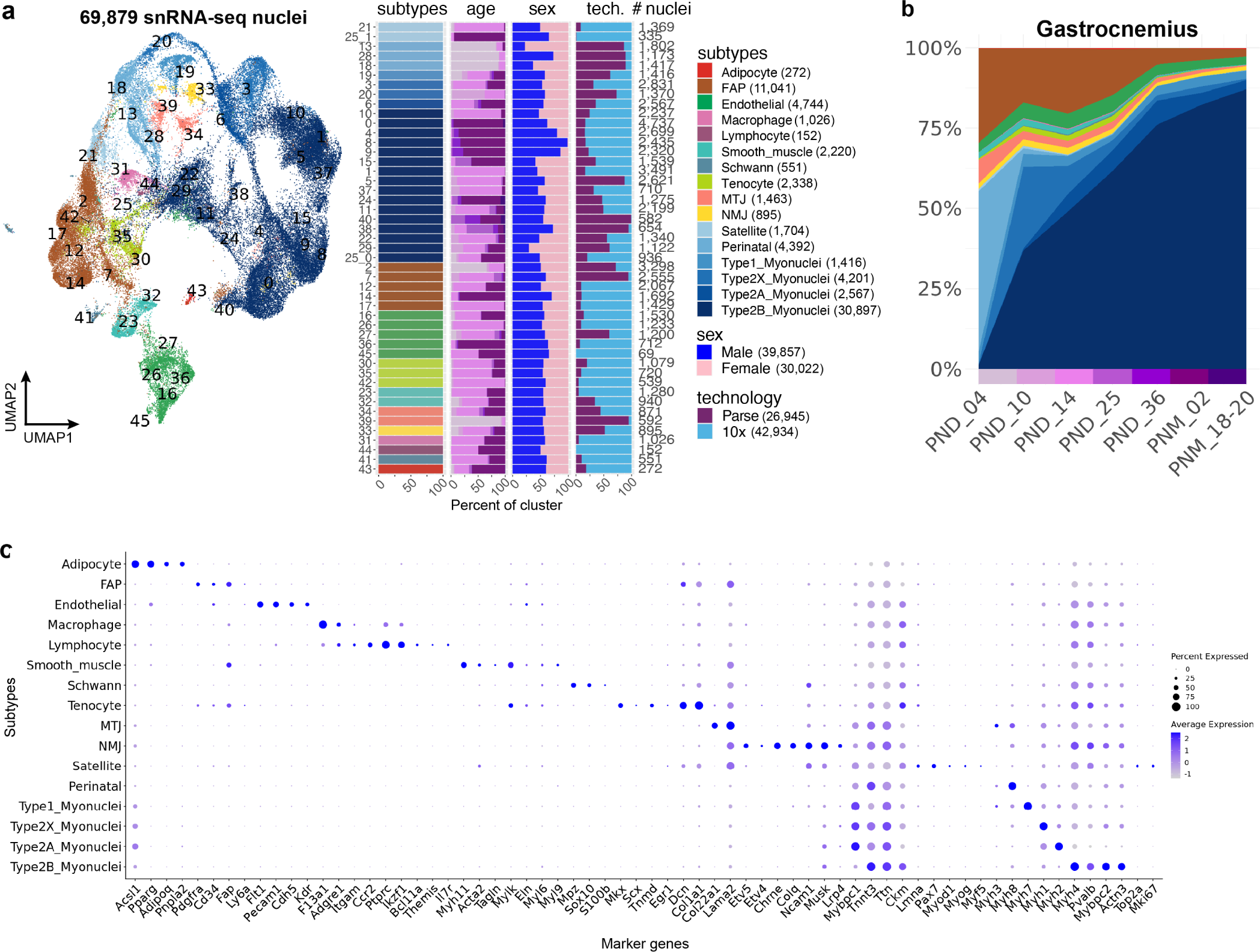
Clustering and annotation of integrated gastrocnemius snRNA-seq data. **a,** UMAP representation of 69,879 gastrocnemius nuclei integrated between Parse and 10x Multiome platforms and breakdown of age, sex, and technology per cluster. Numbers of nuclei per cluster are annotated to the right of the bar plots, and numbers of nuclei per annotated cell subtype are included in the legend. **b,** Dynamics of cell subtype composition across postnatal development in gastrocnemius, with the same color legend as in a. For consistent sampling at each timepoint, only Parse data is shown. **c,** Expression of marker genes across subtypes in gastrocnemius.

**Figure S6.**
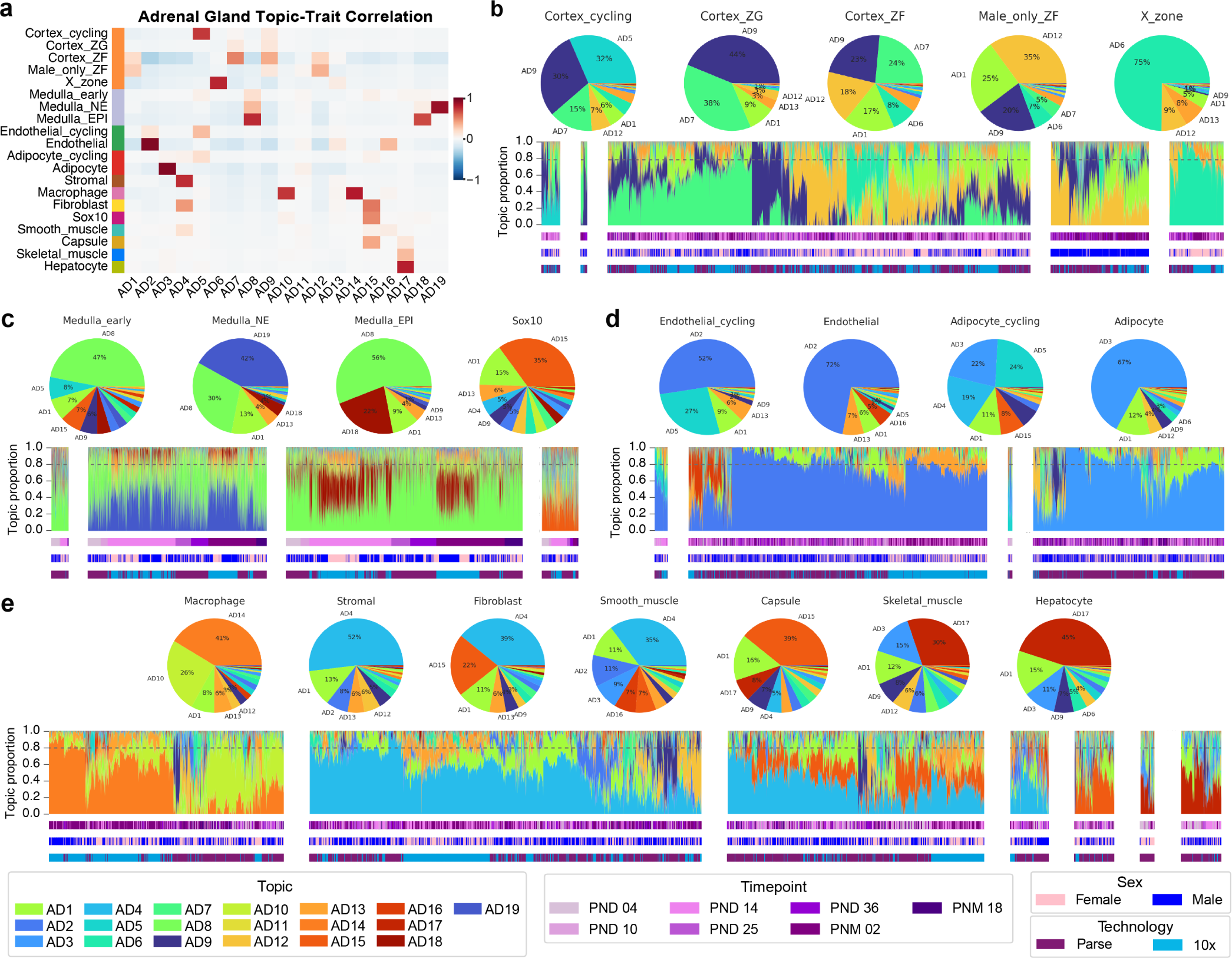
Regulatory topic enrichment and proportions in adrenal gland cell subtypes. **a,** Topic-trait correlation in 19 regulatory adrenal topics. **b,** Structure plots in adrenal cell subtypes, summarized in above pie charts. Topics AD7, AD9, AD12, and AD6 are specific to adrenal cortex. **c,** AD19, AD8, and AD18 are specific to adrenal medulla, while AD15 is specific to *Sox10*+ progenitor cells. **d,** AD2 is endothelial-specific and AD3 is adipocyte-specific. AD5 is a general cycling topic enriched in proliferating cells regardless of subtype. **e,** Topics AD14 and AD10 are specific to macrophages, and topic AD4 is shared across stromal, fibroblast, and smooth muscle cells. AD15 is enriched in the adrenal capsule and fibroblasts.

**Figure S7.**
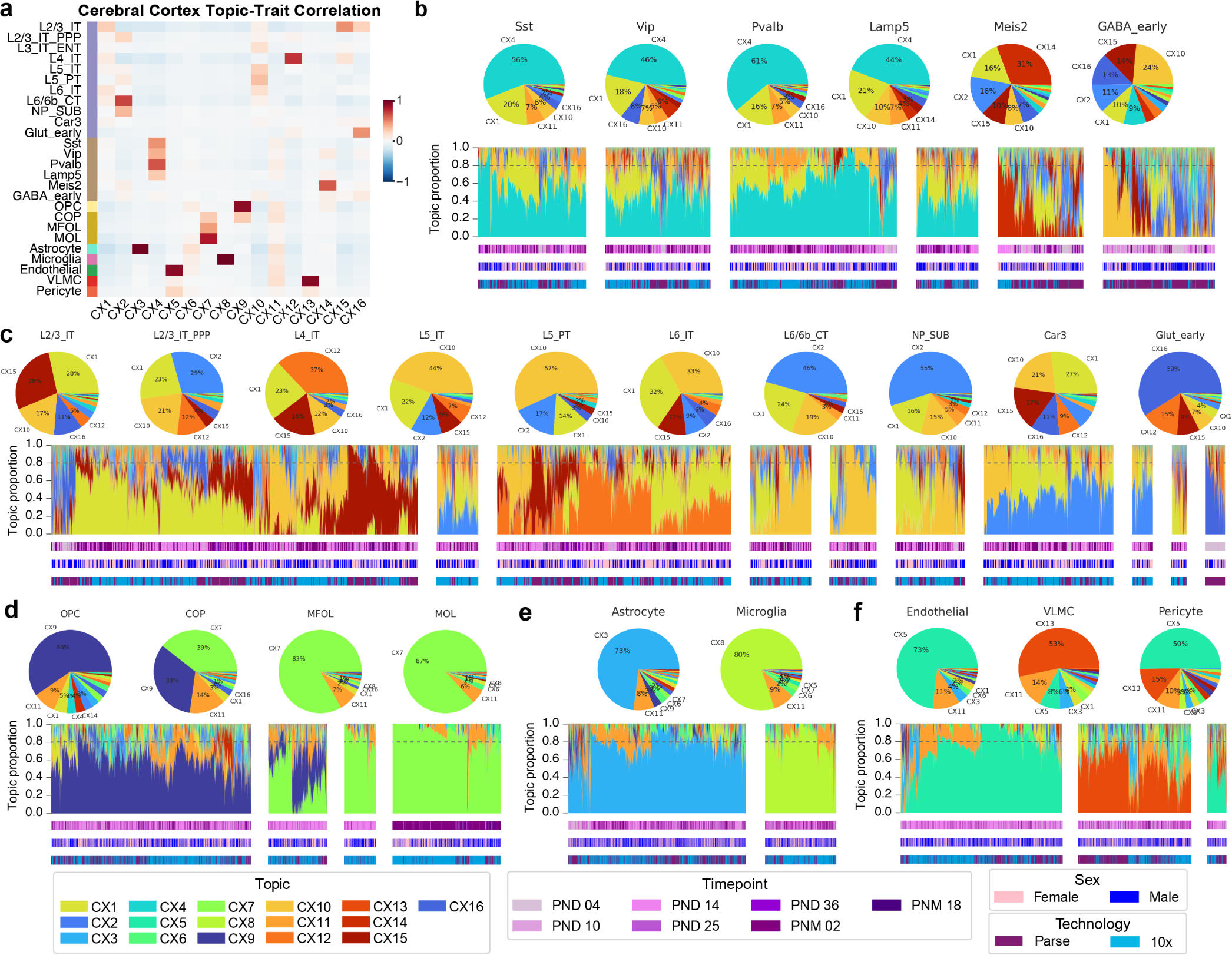
Regulatory topic enrichment and proportions in left cerebral cortex cell subtypes. **a,** Topic-trait correlation in 16 regulatory cortex topics. **b,** Structure plots in cortex cell subtypes, summarized in above pie charts. CX4 is a general GABAergic topic other than *Meis2*+ and early GABAergic cells, which are described by a mix of topics. **c,** Topics CX1, CX2, CX10, and CX12 are all enriched in various excitatory neuronal subtypes. **d,** CX9 is enriched in OPC and COP progenitors, while CX7 is enriched in mature oligodendrocytes. **e,** CX3 is astrocyte-specific and CX8 is microglia-specific. **f,** CX5 is enriched in endothelial and pericytes and CX13 is specific to VLMC (vascular leptomeningeal cells).

**Figure S8.**
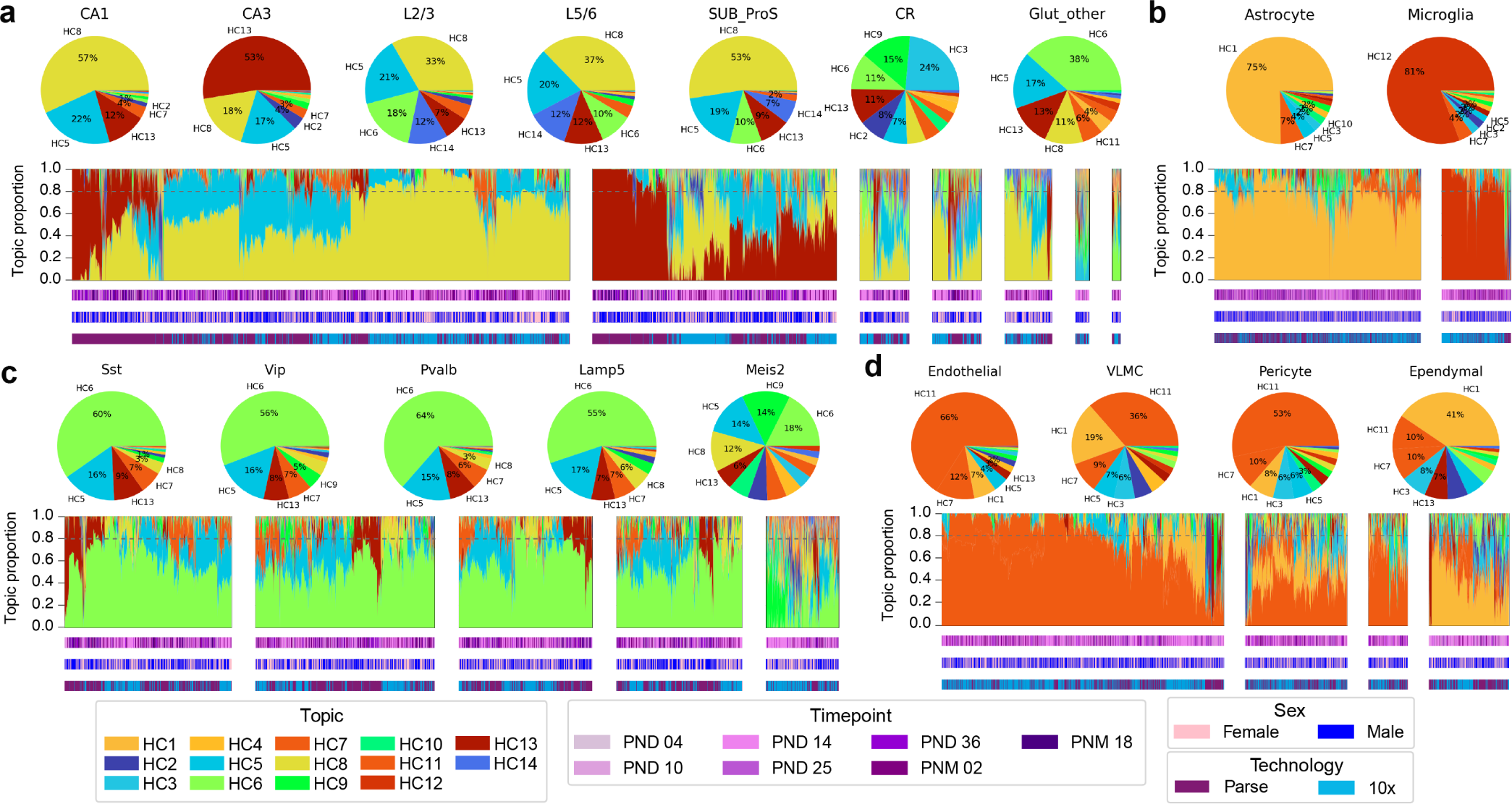
Regulatory topic proportions in left hippocampus cell subtypes. **a,** Structure plots in hippocampus cell subtypes, summarized in above pie charts. HC8 is enriched in CA1 and shared across various other glutamatergic subtypes, and HC13 is CA3-specific. **b,** HC1 is astrocyte-specific, while HC12 is microglia-specific. **c,** HC6 and HC5 are general GABAergic neuron topics, while the *Meis2*+ subtype is described by a mix of topics. **d,** HC11 is enriched in endothelial, pericytes, and VLMC (vascular leptomeningeal cells), while HC1 is shared in VLMC and ependymal cells.

**Figure S9.**
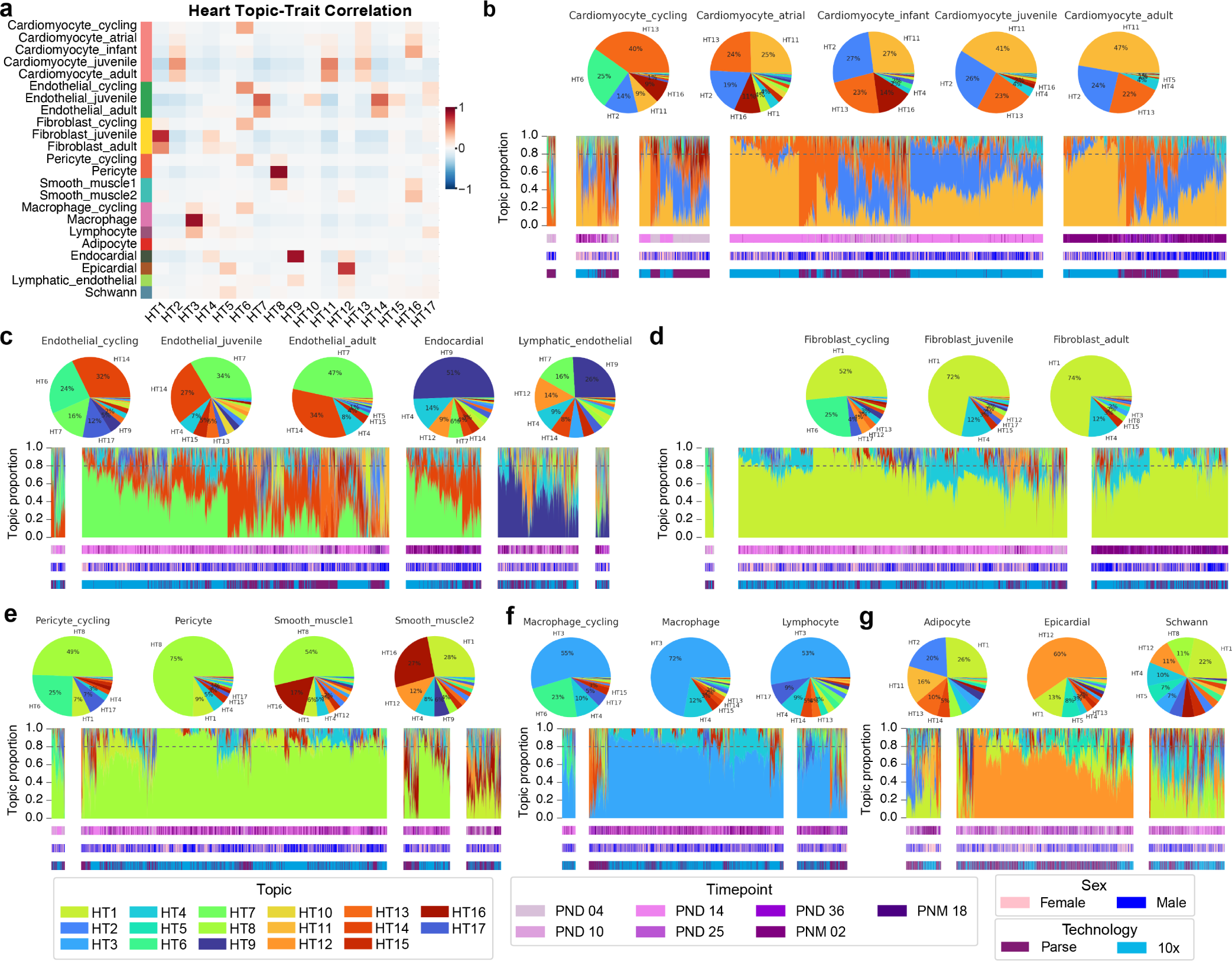
Regulatory topic enrichment and proportions in heart subtypes. **a,** Topic-trait correlation in 17 regulatory heart topics. **b,** Structure plots in heart cell subtypes, summarized in above pie charts. Topics HT2, HT11, and HT13 are shared by cardiomyocytes at all developmental stages. **c,** HT7 and HT14 are enriched in endothelial cells, while HT9 is enriched in endocardial and lymphatic endothelial cells. **d,** HT1 is enriched in cardiac fibroblasts at all developmental stages. **e,** HT8 is specific to pericytes and one subtype of smooth muscle, while the other smooth muscle subtype is enriched in HT16 and HT1. **f,**HT3 is the macrophage-specific topic in heart. HT6 is a general cycling topic enriched in proliferating cells regardless of subtype. **g,** HT12 is specific to epicardial cells. Adipocytes and Schwann cells are made up of several topics, the largest fraction being HT1 which is also shared with fibroblasts and smooth muscle.

**Figure S10.**
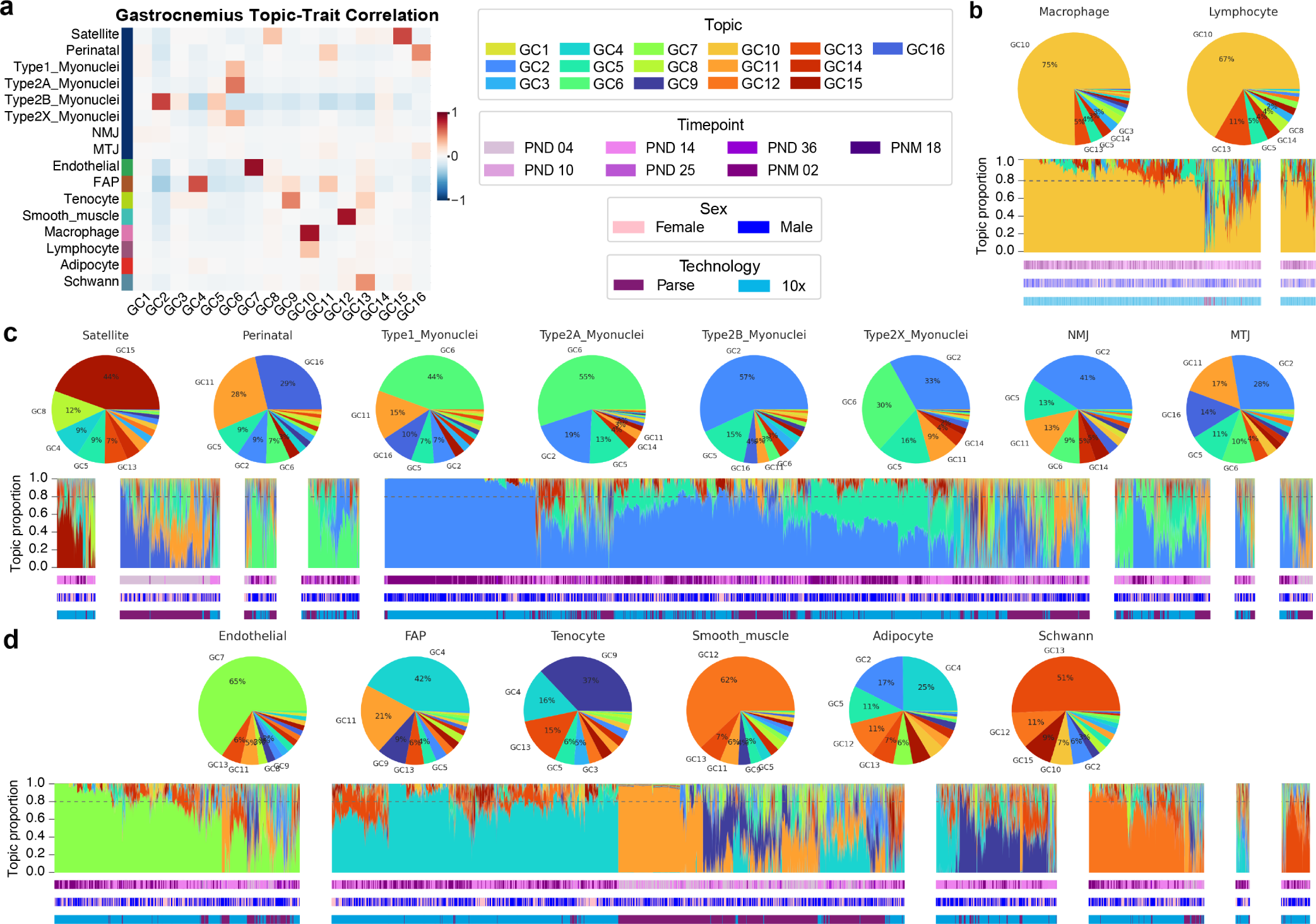
Regulatory topic enrichment and proportions in gastrocnemius subtypes. **a,** Topic-trait correlation in 16 regulatory gastrocnemius topics. **b,** Structure plots in gastrocnemius cell subtypes, summarized in above pie charts. GC10 is enriched in both macrophages and lymphocytes. **c,** Topics GC2, GC5, GC6, and GC11 are shared across mature myofiber subtypes. Most cell participation in type 2B and type 2X is attributed to topic GC2, but type 2X also shares GC6 with type 2A and type 1. Perinatal myonuclei are described by GC16 and GC11, while GC15 and GC8 are specific to satellite cells. Specialized NMJ (neuromuscular junction) and MTJ (myotendinous junction) myonuclei have no specific regulatory topic, but share a mix of muscle-enriched topics. **d,** GC7, GC12, and GC13 are specific to endothelial, smooth muscle, and Schwann subtypes, respectively. FAP (fibro-adipogenic progenitors) are enriched for GC4 and GC11 which are also timepoint-specific, with GC11 enriched in infants and GC4 specific to adults and juveniles.

**Figure S11.**
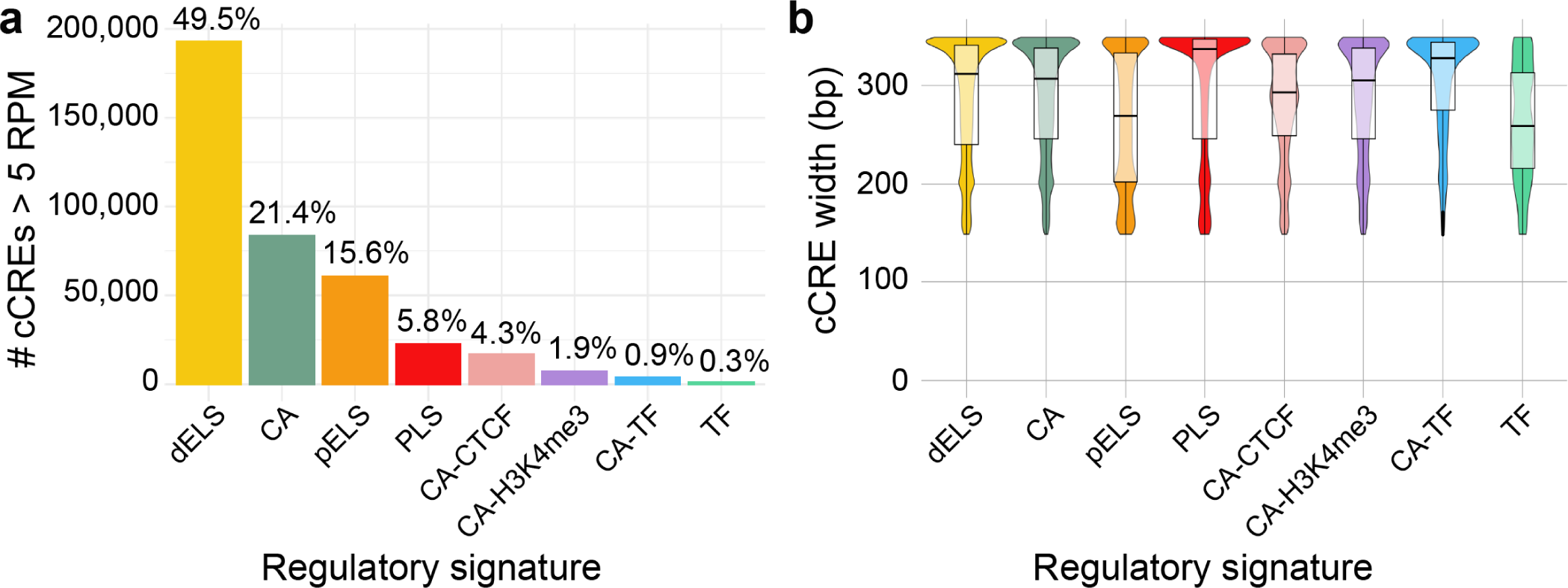
cCRE classification by regulatory signature. **a,** Breakdown of 390,146 cCREs >5 RPM in at least 1 pseudobulk cluster in 10x Multiome snATAC-seq data across all 5 tissues. Most cCREs are classified as dELS (distal enhancer-like signature), CA (chromatin accessible), and pELS (proximal enhancer-like signature). Less than 15% of accessible cCREs are CA-CTCF (chromatin-accessible CTCF), CA-H3K4me3 (chromatin-accessible with promoter-associated histone modification), CA-TF (chromatin-accessible, TF signal), and TF (TF signal). **b,** All cCREs are between 150 and 350 bp with an average of 284 bp with consistent distributions across the 8 categories. Therefore, we opted to normalize snATAC-seq quantifications across the cCREs using reads-per-million (RPM).

**Figure S12.**
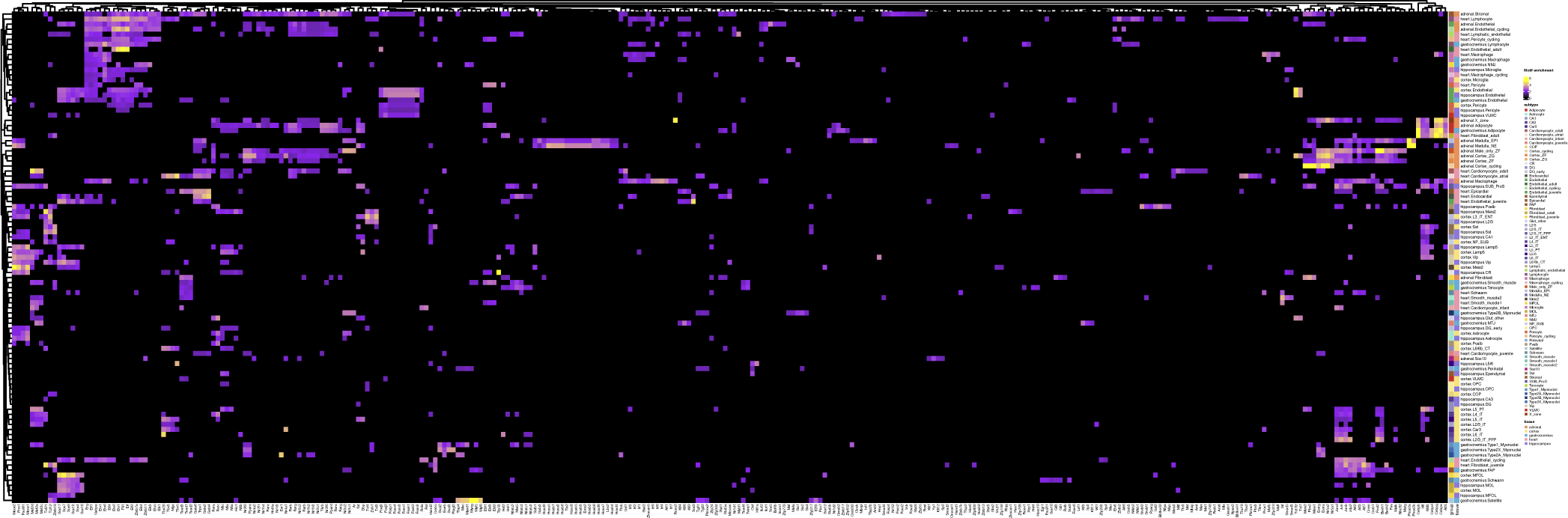
Motif enrichment in subtype-specific cCREs across all tissues. Out of 765 possible JASPAR motifs, 317 were enriched in at least 1 subtype with an adjusted p-value *≤* 0.05, enrichment *≥* 1.5, and bulk RNA-seq expression *≥* 5 TPM in at least 1 sample in the tissue.

